# Mitochondrial Integrated Stress Response Activation Creates a Therapeutic Vulnerability to MCL-1 Inhibition in Acute Myeloid Leukemia

**DOI:** 10.64898/2025.12.01.691686

**Authors:** Lauren Brakefield-Laird, Amit Budhraja, Peter M. Hall, Dixie Brewington, Jamila Moore, Josi Lott, Yonghui Ni, Dennis Voronin, Oliver Grant-Chapman, Tresor Mukiza, Tristen Wright, Yong-Dong Wang, Sandi Radko-Juettner, Shondra M. Pruett-Miller, Stanley Pounds, Peter Vogel, Joseph T. Opferman

**Author notes:** Correspondence should be addressed to Joseph T. Opferman.: St. Jude Children’s Research Hospital, MS#340, 262 Danny Thomas Place, Memphis, TN 38105, USA.

## Abstract

MCL-1 (myeloid cell leukemia-1) promotes survival and confers therapeutic resistance in acute myeloid leukemia (AML), particularly in high-risk subtypes harboring *KMT2A* rearrangements (*KMT2A*-r). Clinical trials of patients with hematological malignancies treated with MCL-1 inhibitor monotherapy have revealed dose-limiting toxicity and poor response rates. Therefore, we sought to identify combinatorial treatment approaches to enhance the efficacy of MCL-1 inhibitors with the goal of improving response rates and limiting toxicities. Here, we report the inhibition of electron transport chain (ETC) complex I (CI) function as a synthetic lethal partner for MCL-1 inhibition. Co-targeting CI and MCL-1 synergistically reduces the viability in AML cell lines and patient-derived xenograft (PDX) samples *in vitro*, while significantly prolonging survival in mice bearing PDX AML, indicating the preclinical potential for combinatorial therapy. These findings provide a mechanistic rationale and preclinical evidence for dual inhibition of MCL-1 and CI as a therapeutic strategy, offering a potential path to overcome resistance to single-agent MCL-1 inhibitors and improve outcomes for patients with high-risk AML. Mechanistically, we reveal that CI inhibition induces the activation of the integrated stress response (ISR), resulting in ATF4 activation downstream of the eIF2α kinase, HRI. The activation of HRI by CI inhibition is dependent on the mitochondrial stress messenger, DELE1. Together, these results indicate that co-inhibition of MCL-1 and ETC CI function has the potential for improving responses in patients with *KMT2A*-r AML.

## Introduction

Acute myeloid leukemia (AML) is an aggressive hematologic malignancy characterized by clonal expansion of myeloid progenitor cells and impaired differentiation (ref. 1). Clinical outcomes vary widely, with certain genetic subtypes, including *KMT2A* (previously known as *MLL*) rearrangements (hereafter referred to as *KMT2A*-r), representing high-risk disease associated with poor prognosis and resistance to standard therapies (ref. 1, 2). The intrinsic pathway of apoptosis is regulated by the BCL-2 family of proteins, where anti-apoptotic members such as BCL-2 and MCL-1 promote survival, and pro-apoptotic effectors like BAX and BAK drive cell death (ref. 3). Dysregulation of this balance contributes to therapeutic resistance, making BCL-2 family proteins attractive therapeutic targets in cancer. BH3-mimetics are small molecules that mimic the function of pro-apoptotic BH3-only proteins to inhibit anti-apoptotic BCL-2 family members and induce apoptosis in cancer cells (ref. 4). Venetoclax is a selective BCL-2-targeting BH3-mimetic that is approved for use in combination with hypomethylating agents or low-dose cytarabine for the treatment of newly diagnosed AML in patients who are ineligible for intensive chemotherapy (ref. 5, 6). However, resistance often emerges through compensatory upregulation of MCL-1 (ref. 7, 8, 9, 10, 11, 12). MCL-1 inhibitors have demonstrated potent activity in preclinical AML models and some have been evaluated in clinical trials; however their development has been constrained by observed toxicities (ref. 13, 14, 15, 16).

AML is a heterogeneous disease, not only at the genetic and epigenetic levels but also in its cellular and metabolic composition (ref. 17, 18). Among AML populations, leukemic stem cells (LSCs) display distinct metabolic programs compared with bulk leukemia cells, often relying on mitochondrial oxidative phosphorylation (OXPHOS) rather than glycolysis for energy production and survival (ref. 19, 20). This metabolic dependency contributes to LSC quiescence, resistance to conventional chemotherapies, and ability to drive disease relapse (ref. 21, 22). The metabolic heterogeneity of AML therefore represents both a challenge and an opportunity: by identifying and targeting specific metabolic vulnerabilities in LSCs, such as OXPHOS or mitochondrial stress pathways, it may be possible to selectively eradicate these therapy-resistant cells without harming normal hematopoietic stem cells. Targeting the unique metabolic dependencies of LSCs offers a promising strategy to enhance therapy, prevent relapse, and eradicate AML at its root.

Targeting mitochondrial metabolism, particularly electron transport chain (ETC) complex I (CI), has emerged as a promising strategy in hematologic malignancies, including AML. CI is the first enzyme complex of the mitochondrial electron transport chain and is important for facilitating OXPHOS and ATP production (ref. 23). Inhibiting CI disrupts mitochondrial respiration and induces mitochondrial stress (ref. 24). IACS-010759 (hereafter referred to as IACS) is a potent, selective small-molecule CI inhibitor that has shown preclinical efficacy in AML models (ref. 25, 26, 27). Other experimental CI inhibitors are being investigated with similar mechanistic rationales, emphasizing the vulnerability of metabolically dependent cancer cells (ref. 28, 29, 30, 31). Clinical evaluation of IACS demonstrated activity in relapsed/refractory AML and other malignancies, although dose-limiting toxicities, including lactic acidosis and neurotoxicity, have limited its use as monotherapy (ref. 25). Preclinical studies suggest that combining CI inhibitors with other therapies has the potential to selectively eradicate LSCs and overcome resistance, highlighting CI inhibition as a promising avenue to exploit AML’s metabolic vulnerabilities and improve long-term treatment outcomes (ref. 32, 33, 34, 35, 36).

The mitochondrial stress response, particularly the DELE1-HRI-ATF4 axis, is a critical mechanism by which cells sense and adapt to mitochondrial dysfunction. Upon stress, DELE1 can be cleaved, which then activates the kinase HRI to phosphorylate eIF2α, leading to induction of the transcription factor ATF4 (ref. 37, 38). ATF4 can then induce the expression of genes controlling stress adaptation, metabolism, and apoptosis (ref. 39, 40, 41, 42). In mouse models of AML, the repression of ATF4 impairs AML growth, delaying leukemogenesis and reducing leukemia-initiating cell activity (ref. 43). These findings underscore the therapeutic potential of the mitochondrial stress response: targeting DELE1, HRI, or ATF4 could selectively disrupt AML cell adaptation to stress, enhance sensitivity to treatment, and improve outcomes, making this pathway a promising target for novel AML therapies.

In this study, we investigated how metabolic modulation can enhance the efficacy of MCL-1 inhibition in AML. We identified a synthetic lethal interaction between inhibition of mitochondrial CI and MCL-1 blockade. Specifically, the selective CI inhibitor IACS synergized with multiple MCL-1 inhibitors to enhance AML cell death. Mechanistically, this combination activated the DELE1-HRI-ATF4 arm of the integrated stress response (ISR), driving cell death. These results highlight the therapeutic value of co-targeting MCL-1 and mitochondrial metabolism to achieve robust anti-leukemic activity in both *in vitro* and *in vivo* models.

## Materials and Methods

### Cells and cell culture

MV4-11 cells were acquired from the American Type Culture Collection (ATCC; Manassas, VA, USA). ML2, THP1, OCI-AML3, U937, HL60, CHRF-288-11, M-O7E, and Molm-13 cells were a gift from Charles Mullighan at St. Jude Children’s Research Hospital (SJCRH, Memphis, TN, USA). AML cell lines were cultured in Roswell Park Memorial Institute 1640 media (Thermo Fisher Scientific, Waltham, MA, USA) supplemented with 10% fetal bovine serum (Gibco, Billings, MT, USA), 55 µM 2-mercaptoethanol, 2 mM glutamine, and 1% penicillin-streptomycin solution (Invitrogen, Waltham, MA, USA). Cell lines were authenticated via short tandem repeat profiling through the Hartwell Center for Biotechnology (SJCRH; Memphis, TN, USA). Cas9-expressing MV4-11 cells were generated using lentiCas9-Blast, a gift from Feng Zhang (Addgene viral prep# 52962-LV). Stably expressing single cell clones were generated under blastocidin selection (15 μg/mL). All cell lines were subjected to frequent mycoplasma testing using the MycoAlert Mycoplasma Detection Kit (Lonza, Basel, Switzerland) by following the manufacturer’s protocol and were maintained in culture for no more than 6 weeks. S63845 (Cat. No. HY-100741), S64315 (Cat. No. HY-112218), and IACS-010759 (Cat. No. HY- 112037) were obtained through MedChem Express (Monmouth Junction, NJ, USA). AMG 176 and AZD5991 compounds were obtained through the Department of Chemical Biology and Therapeutics (SJCRH). Thapsigargin was obtained through Millipore Sigma (Burlington, VT, USA). A-1331852 was obtained through Medkoo Biosciences (Durham, NC, USA), venetoclax (ABT-199) was from LC Laboratories (Woburn, MA, USA), and navitoclax (ABT-263) was from ApexBio Technology LLC. (Houston, TX, USA).

### *Ex Vivo* patient-derived xenograft isolation and culture

Leukemia cells originating from two pediatric patients (SJAML016500_D1 and SJAML030440_R2) harboring *KMT2A*-r AML were established at St. Jude Children’s Research Hospital and can be accessed using the Public Resource of Patient-derived and Expanded Leukemia (PROPEL) repository (https://propel.stjude.cloud/). SJAML016500_D1 was derived from a 12-year-old patient at the time of diagnosis, and SJAML030440_R2 was derived from a 5-year-old patient at relapsed disease and labeled in figures respectively as “diagnosis” and “relapsed”. For *in vivo* propagation, 1.5–1.75E6 cells were administered by tail vein injection into immunodeficient Non-Obese Diabetic (NOD) Cg-*Prkdc^scid^ Il2rg^tm1Wjl^* Tg(CMV-IL3, CSF2, KITLG)1Eav/MloySzJ (NSG-SGM3) 8- to 12-week-old female mice (Jackson Laboratories, Bar Harbor, ME, USA). Retro-orbital bleeds were performed every 4 weeks until a humane endpoint (presence of hind leg weakness/paresis, delayed response to touch, hunched posture, tachypneic) was reached. Disease progression was assessed by flow cytometry using a panel to detect mouse CD45, human CD19, human CD34, human CD33, human CD7, human CD45, and DAPI. Once euthanized, bone marrow and spleen cells were harvested by flushing the femurs or mashing the spleens through a 100 μm filter. Collected cells were centrifuged at 1200 RPM for 5 min at 4° C and underwent red blood cell lysis using the BD Pharm Lyse lysing reagent (BD Biosciences, Franklin Lakes, NJ, USA) using the manufacturer’s protocol. Mouse cells were depleted using mCD45-biotin antibody clone 30-F11 (Cat # 103104, Biolegend San Diego, CA, USA), Streptavidin magnetic particles (Roche, Basel, Switzerland), and a Miltenyi magnet system based on the manufacturer’s protocol. *Ex vivo* cells were maintained for up to 24 h in StemSpan Serum-Free Expansion Medium II (StemCell Technologies, Vancouver, BC, Canada) supplemented with 10 ng/mL GCSF, 100ng/mL SCF, 10 ng/mL IL3, 100 ng/mL IL6, 100 ng/mL TPO, and 100 ng/mL FLT3 ligand (Peprotech, Cranbury, NJ, USA).

### CRISPR knockout library design, assembly, and validation

The ISR CRISPR knockout KO library was designed using the CRISPick tool and composed of up to five high-confidence *Streptococcus pyogenes* single-gRNAs (sgRNAs) per gene (ref. 44, 45) by the Center for Advanced Genome Engineering Core Facility (CAGE, SJCRH). Briefly, CRISPick was used to identify the top candidate sgRNAs for each gene, which were then subjected to another round of in-house off-target filtering. Additionally, non-targeting control (NTC) sgRNAs were incorporated to comprise approximately 10% of the final library composition. Oligos with appropriate overhangs were generated (TWIST Bioscience, South San Francisco, CA, USA). Oligo library amplification and Gibson Assembly into the lentiGuide Puro vector backbone (Addgene #52963) was performed as previously described (ref. 46, 47). The ISR KO plasmid library underwent assembly, amplification, and also validation in the Center for Advanced Genome Engineering following the Broad Institute GPP protocol (https://portals.broadinstitute.org/gpp/public/resources/protocols) with the only modification being the use of Endura DUOs electrocompetent cells. The Single-end 100-cycle sequencing was performed on a NovaSeq 6000 (Illumina, San Diego, CA, USA) by the Hartwell Center for Genome Sequencing Facility at SJCRH. The Human CRISPR Metabolic Gene Knockout Library DNA was a gift from David Sabatini (Addgene #110066) and was amplified in the same manner (ref. 48). Validation of both libraries were performed to assess for sgRNA presence and representation using calc_auc_v1.1.py (https://github.com/mhegde/) and count_spacers.py (ref. 46). A list of genes and sgRNA sequences for the ISR KO library is available in Supplementary Table S1.

### CRISPR KO library lentiviral particle production

Lentiviral particles for the Human CRISPR Metabolic KO Library and ISR KO library were manufactured by the Vector Development and Production laboratory (SJCRH) as previously described (ref. 49). Briefly, 2e6 SJ293TS-DPB cells per mL (500mL total) were transfected with either the Human CRISPR Metabolic KO or ISR KO library plasmid DNA and the helper plasmids, pCAG-kGP1-1R-AF (gagpol), pCAG-VSVG-AF (VSV-G), and pCMV-RTR2-AF (Rev), at a ratio of 14:4:2:0.25, respectively. Six hours after transfection, cells were diluted in 500 mL with freestyle culture media and treated with Benzonase at a final concentration of 6.25 U/mL. After 48 h, vector supernatants were harvested, clarified by centrifugation, and filtered through both 0.45- and 0.22- µm filters. The lentiviral containing supernatant was adjusted to 300 mM NaCl with 50 mM Tris pH 8.0 before being loaded onto Mustang Q XT5 ion-exchange capsule (Cytiva, Marlborough, MA) per the manufacturer’s instructions using an Akta Avant chromatography system (Cytiva). Lentiviral particles were eluted from the column by using 2 M NaCl, 50 mM Tris pH 8.0. Eluates from the Mustang Q XT5 ion-exchange capsule were diluted 5-fold with phosphate buffered saline (PBS) and diafiltrated twice in PBS using a Vivaflow 50 with a 100,000 molecular weight cutoff (Sartorius AG, Goettingen, Germany) according to the manufacturer’s instructions. The final diafiltration was performed in X-VIVO 15 media (Lonza). The purified and concentrated vectors were passed through a 0.22- µm filter and stored at -80° C. To determine viral titers, purified/concentrated vector preps were titered on HOS-6A5 cells. Transduced HOS6A5 cells were harvested 4 days post-transduction. Droplet digital PCR was used to determine the viral titer as follows: genomic DNA was isolated from transduced cells, digested with MspI, and used as a template in PCR with a probe against the HIV Psi element of the vector and a probe against human RPP30, and endogenous control assumed to be ∼2 copies/cell. The primer-probe sets listed in Supplementary Table S2 were used to amplify the endogenous control gene, *RPP30* and the HIV psi sequence.

### CRISPR KO screens

The Human CRISPR Metabolic Gene KO Library was a gift from David Sabatini (Addgene #110066) (ref. 48). MV4-11 cells were transduced with the Human CRISPR Metabolic Gene KO Library at a multiplicity of infection (MOI) of 0.5 with a sgRNA coverage of 1000x. Twenty-four hours after the transduction, 2 µg/mL of puromycin was added for selection and kept in the media for 7 d and then removed from the media. Cells were then cultured in media supplemented with 10 nM S64315, 350 nM AMG 176, 90 nM AZD5991, or volume-matched DMSO for another 24 h, followed by culture in drug-free media for up to 14 population doublings. MV4-11 Cas9 cells were transduced with the ISR KO library at an MOI of 0.5 and a sgRNA coverage of 5000x. Twenty-four hours after transduction, puromycin was added for selection in the same manner as previously mentioned. Cells were cultured in media containing DMSO or a combination of 4 nM S64315 and 160nM IACS. The drugs were washed from the media after 24 h of drug exposure and the cells were cultured for 13 population doublings in drug-free media. For both screens, genomic DNA was extracted from the surviving cells using DNeasy Blood and Tissue Kit (Qiagen, Hilden, Germany) following the manufacturer’s instructions, then amplified by PCR for sequencing of sgRNAs via Nextseq (Illumina). The MAGeCK-VISPR algorithm was used to analyze the sequencing reads (ref. 50). Top hits were loaded into Enrichr for pathway analyses (ref. 51, 52).

### KO cell line creation

MV4-11 and THP1 gene KO cell pools were generated using CRISPR-Cas9 technology in the Center for Advanced Genome Engineering (CAGE, SJCRH). Two million cells were transiently transfected with precomplexed ribonuclear protein containing 300 pmol of each chemically modified sgRNA (Synthego, Redwood City, CA, USA) and 100 pmol 3X NLS SpCas9 protein (SJCRH Protein Production Core) via nucleofection (Lonza 4D-Nucleofector™ X-unit) using solution SF and program DJ-100 in a 100-µL cuvette according to the manufacturer’s protocol. Three days after nucleofection, cell pellets with approximately 100,000 cells were lysed and gene-specific amplicons were generated and then sequenced by targeted next-generation sequencing as previously described (ref. 53). Next-generation sequencing data were processed using CRIS.py (ref. 54). After expansion, the cell pools were authenticated using the Power Plex® Fusion System (Promega, Madison, WI, USA) by the Hartwell Center for Biotechnology (SJCRH) and tested for mycoplasma by the MycoAlert^TM^Plus Mycoplasma Detection Kit (Lonza). KO pools were cloned out to achieve single-cell KO clones that were then further verified by western blot and next-generation sequencing. Editing construct and NGS primer sequences are listed in Supplementary Table S3.

### Immunoblotting and antibodies

Protein was extracted by resuspension of cell pellets in RIPA buffer supplemented with final concentrations of 10 mM sodium fluoride (MilliporeSigma Burlington, MA, USA) 1 mM sodium orthovanadate (MP Biomedicals, Santa Ana, CA, USA), 1x protease inhibitor (MilliporeSigma), and 1 mM phenylmethylsulfonylfluoride (Thermo Fisher Scientific). Gels were loaded with 30-40 μg of protein, run on NuPage 4%-12% bis-tris gradient gels (Invitrogen), and transferred to Immobilon PVDF membranes (MilliporeSigma). The antibodies used were anti-ATF4 (Cl. D4B8 Cat No: 11815S), anti-BAK (Cat No: 3814S) anti-BAX (Cat. No: 2772S) from Cell Signaling Technologies (Danvers, MA, USA), anti-EIF2AK1 (Cat No: 20499-1-AP) from Proteintech (Rosemont, IL, USA) and anti-Actin (Cl. C4. Cat No: MAB1501) from MilliporeSigma. Anti-rabbit and anti-mouse horseradish peroxidase-conjugated secondary antibodies were obtained from Invitrogen. Immunoblots were developed using Odyssey Fc from LI-COR Biosciences (Lincoln, NE, USA), and Image Studio v6.0.0.28 was used to produce the western blot figures. All experiments are representative of three, or more, independently performed experiments.

### Cell death experiments

Cells were seeded in 96-well plates between 50,000 to 100,000 cells per well and treated with the indicated drugs (IACS, AMG 176, AZD5991, S63845, S64315, A-1331852, venetoclax, or navitoclax solubilized in DMSO, or DMSO–vehicle control) for 24 h unless otherwise indicated. For the generation of dose response curves, the cells were treated with BH3 mimetics over a concentration range of 40 µM - 40 nM in 2-fold serial dilutions. For combination treatments, IACS was combined with the MCL-1 inhibitors at the concentrations indicated on the figures.

### Flow cytometry analysis

Flow cytometry data were collected using BD FACS Canto II (BD Biosciences) and data were analyzed using FlowJo Software (v10.10). For cell death experiments, APC Annexin V (Biolegend) was used at a concentration of 1:100. Propidium iodide (MiliporeSigma) was used at a final concentration of 0.5 μg/ mL. Cell viability data were determined and reported using annexin V and propidium iodide double-negative cells as previously described (ref. 55). For dynamic BH3 profiling, the cells were stained over night at 4° C 1:40 with anti-cytochrome *c* antibody (clone 6H2.B4, Biolegend). Flow analysis was performed using previously described gating strategies (ref. 56). The Center of Excellence for Leukemia Studies (CELS) Flow Cytometry Core (SJCRH) collected flow data using the BD FACSymphony A5 (BD Biosciences) to identify and determine the percentage of human CD45^+^ cells as previously described (ref. 57). Each experiment was tested in at least three biological replicates and plotted in GraphPad Prism (v10.5.0) for visualization.

### Quantification of drug synergy

The drug combination responses were calculated based on the Bliss reference model using SynergyFinder+ (ref. 58). The SynergyFinder+ online software and R package were used to generate Bliss scores with p-values from cell viability data. According to the SynergyFinder+ paper, the Bliss p-value is “an empirical p-value to assess the significance of the difference between the estimated average synergy score over the whole dose matrix and the expected synergy score of zero under the null hypothesis of non-interaction” (ref. 58). Scores are considered synergistic if >10 and additive if between -10 to 10. To determine statistical significance between wild-type and knockout genotypes, we obtained the Bliss score and Bliss p-value, then converted the Bliss p-values into Z-scores to calculate the standard deviation of bootstrapping. Finally, we applied a Z-test to compare the difference between the two Bliss scores and obtain a Zp-value. Zp-value less than 0.05 are considered statistically significant.

### ATP production

ATP production was measured using the Seahorse XF Mito Stress Test (Agilent, Santa Clara, CA, USA) and was performed following the manufacturer’s protocol on a Seahorse XFe96 Analyzer. Briefly, cell lines and PDX cells were treated with DMSO or 50 nM IACS in the culture media for 1 h and then plated at a density of 100,000 cells per well into 96-well Seahorse plates that were precoated with 12 μg/mL Cell Tak prepared according to the manufacturer’s protocol (Thermo Fisher Scientific). Before plating, culture media was exchanged for Seahorse XF RPMI Medium supplemented with 1 mM sodium pyruvate, 2 mM L-glutamine and 10 mM glucose. IACS was kept in the media for the duration of the assay. Cells were placed in a CO_2_-free incubator to outgas for 1 h. The Mito Stress Test was performed using a final concentration of 1.5 µM oligomycin (Thermo Fisher Scientific), 2 µM FCCP (Millipore Sigma,), and 0.5 µM rotenone/antimycin A (Sigma Life Sciences/Fisher Scientific). For each biological replicate, 3 or more technical replicates were performed. The total cell number per well was obtained through fluorescent cell counting of cells stained with 15 μM Hoechst 33342 (Thermo Fisher Scientific) on Cytation 5 (BioTek, Winooski, VT, USA) (v 1.0.0-694). Data was normalized to total cell number and analyzed using Agilent Seahorse Analytics online software v1.0.0-694. Mitochondrial and glycolytic ATP rates were calculated using the Agilent Seahorse Analytics online software, and graphs were made in GraphPad Prism (v10.5.0) (ref. 59). Each experiment was performed n=3 or more times. Graphs in the figures depict representative experiments.

### BH3 profiling

Dynamic BH3 profiling was conducted using a previously published method (ref. 60). Briefly, 500,000 cells were seeded and treated with either DMSO or 50nM IACS for 24 h. Cells were collected, washed with PBS and stained with AquaZombie viability dye (Biolegend). Cells were exposed to a 0.0001% digitonin (Thermo Fisher Scientific) for membrane permeabilization and then exposed to increasing concentrations up to 10 μM of the human BIM peptide (sequence: MRPEIWIAQELRRIGDEFNA) for 1 h at 23° C. After BIM peptide exposure, 4% formaldehyde in PBS (Thermo Fisher Scientific) solution was added to terminate the peptide exposure, followed by neutralization with N2 buffer (1.7M Tris base, 1.25M glycine, pH 9,1). The cells were stained with 1:40 with anti-cytochrome *c* antibody (clone 6H2.B4, Biolegend) and stained overnight. The next day, cells were washed twice with 2% FBS in PBS and cytochrome c release was assessed by flow cytometry.

### RNA sequencing

Five million AML cells were plated and treated with DMSO or 50 nM IACS-010759 for 8 or 24 h for three independent replicates. Total RNA was isolated using the RNeasy kit (Qiagen) following the manufacturer’s protocol with the on-column DNA digest step included. RNA sequencing libraries for each sample were prepared with 1 μg total RNA by using the Illumina TruSeq RNA Sample Prep v2 Kit per the manufacturer’s instructions, and sequencing was completed on the Illumina NovaSeq 6000. The 100-bp paired-end reads were trimmed, filtered against quality controls (Phred-like Q20 or greater) and length (50 bp or longer), and aligned to the human reference sequence GRCh38 (hg38) using CLC Genomics Workbench v20 (Qiagen). For gene expression comparisons, the transcript per million counts were obtained from the CLC RNA-Seq Analysis tool. Differential gene expression analysis was performed with a non- parametric ANOVA followed by Kruskal-Wallis and Dunn’s *post hoc* tests on log-transformed transcript per million counts between three replicates of experimental groups, implemented in Partek Genomics Suite v7.0 (Partek Inc., San Diego, CA, USA). The significantly differentially expressed genes were identified with p-value < 0.05 and |log2R| > 0.585 (half-fold change).

### *In vivo* animal studies

For *in vivo* combination therapy studies, 8- to 12-week-old, female NSG-SGM3 mice were used and housed under a 12 h–12 h light–dark cycle (light on at 06:00 and off at 18:00), with food and water provided *ad libitum*. Animals were housed in the facility at 20–22° C with humidity levels maintained at 30%–70% at cage level. All procedures were conducted in compliance with protocols approved by the Institutional Animal Care and Use Committee (IACUC) of SJCRH. Mice were injected with 200,000 or 500,000 PDX cells by tail vein injection. Beginning five weeks after injection, peripheral blood was collected weekly from mice via retro-orbital bleeding to assess the presence of human CD45 cells using flow cytometry. Mice meeting enrollment criteria (∼0.5% hCD45+ cells in the peripheral blood) were randomized into treatment arms using a blocked randomization list available from: (https://www.sealedenvelope.com/simple-randomiser/v1/lists). After randomization, no blinding was performed. Mice were treated with vehicles, 5 mg/kg IACS-10759 PO and vehiclefor S64315 IP, 6.5 mg/kg S64315 IP and vehicle for IACS PO, or a combination of 5 mg/kg IACS-10759 + 6.5 mg/kg S64315 once daily, for 5 days on and two days off for two weeks. Two days after the final drug dose, n=3 mice per treatment group were euthanized to assess tumor burden at the end of treatment. The remaining mice were monitored for up to two months post-treatment or until a scientific endpoint (presence of hind leg weakness/paresis, delayed response to touch, hunched posture, tachypneic) was reached for survival analysis. For the survival analysis, the baseline is defined as the start date of treatment. The treatment period lasted 12 days (5 days on, 2 days off, and 5 days on). Mice that reached a humane endpoint and did not receive all drug doses are excluded from the analysis for a total of 17 mice (5 from vehicle, 5 from IACS group, 4 from S64315 group, and 3 from combination).

### *In vivo* formulations

The Analytical Technology Center-Preclinical Formulation Team (ATC-PFT) provided the preclinical formulations. For *in vivo* studies, S64315 free-base was formulated in 2% v/v Kolliphor EL in PBS. IACS-10759 free-base was formulated in 0.5% methylcellulose (type 400 cPs) in UP water. Both formulations were stable for two weeks at room temperature and 4° C at the prepared concentrations.

### Blood serum lactate collections

Blood samples were collected by retro-orbitally and immediately placed in sodium fluoride/potassium oxalate solution at a 1:10 ratio before being placed on ice. Plasma was separated within 15 min of collection by centrifugation at 5600 RPM for 10 min at 4° C. The resulting serum was placed in a separate tube and stored at 4° C overnight. Samples were analyzed using the Point Lactate Reagent Set (Canton, MI, USA) according to the manufacturer’s instructions.

### Data analysis, representation, and statistics

IC_50_ values from single agent dose curves were determined using a nonlinear regression model to fit dose response curves in GraphPad Prism. Flow cytometry data were analyzed using FlowJo. Seahorse data were processed using the online software Agilent Seahorse Analytics (v1.0.0). For survival analysis, the difference in the treatment effect appears more pronounced during the later stages of follow-up, so a weighted log-rank test (Fleming-Harrington test) that emphasizes differences in survival occurring later in time was applied. We used the Fleming-Harrington test for right-censored data based on counting processes (ref. 61), setting the parameters ρ=0 and λ=1 in the weighting function w(t)=[*S*(*t*)]*ρ*[1−*S*(*t*)]*λ*, where S(t) denotes the Kaplan-Meier estimate of survival at time t. The R package ‘FHtest’ was used to calculate this (ref. 62). The resulting p-values from the weighted log-rank test is reported in Figure 5. GraphPad Prism was used for statistical analysis and data visualization. Schematic figures were generated using Biorender. All statistical tests are stated in the figure legends. Statistical significance is represented as follows: P-values or Zp-values ≤ 0.05 are scored as *, ≤ 0.01 are scored as **, ≤ 0.001 are scored as ***, ≤0.0001 are scored as ****. Final figures were made in Adobe Illustrator 2025 (v25.5.1).

## Results

### *KMT2A*-r AML is sensitive to MCL-1 inhibition

To identify a molecular subtype of AML that would benefit from MCL-1 inhibitors, a panel of AML cell lines was treated with increasing doses of various BH3 mimetics to determine the sensitivity of each cell line (Supplementary Fig. S1 and Table 1). The panel includes both pediatric- and adult-derived AML cell lines, different French-American-British classifications, and cytogenetics (ref. 63). The AML cell lines were treated with either vehicle control (DMSO), a BCL-XL inhibitor (A-1331852), MCL-1 inhibitor (S63845), a pan BCL-2, BCL-XL, and BCL-W inhibitor (ABT-263, also known as navitoclax) or a BCL-2 inhibitor (ABT-199, also known as venetoclax). Cell death was assessed by annexin V (AV) and PI-negative cells using flow cytometry after 24 h of treatment. Five AML cell lines were most sensitive to the MCL-1 inhibitor, S63845 (Supplementary Fig. 1A-E and Table 1). This included three of the four AML cell lines (ML2, MV4-11, and THP1) that harbored *KMT2A*-r (Supplementary Fig. S1A-C). In contrast, other AML cell lines responded preferentially to inhibition of BCL-XL (CHRF 288-1) or BCL-2 (HL60) with several cell lines responding preferentially to navitoclax (M-O7E and Molm-13) (Supplementary Fig. S1H and I). *KMT2A*-r occur in up to 60% of infant patients and 15% of pediatric patients with AML and generally reflect an intermediate to poor prognosis in this population (ref. 64, 65, 66). Therefore, we decided to optimize a combination therapy to potentiate responses of *KMT2A*-r AML to cell death induced by MCL-1 inhibition.

**Table 1.** Characterization of sensitivity to BH3 mimetics in AML cell lines. IC_50_ values in nM of the cell lines treated with 40 μM-40 nM of BH3 mimetics for 24 h. The BH3 mimetics used include A-1331852 (BCL-XL inhibitor), S63845 (MCL-1 inhibitor), ABT-263/navitoclax (pan BCL-2, BCL-XL, and W inhibitor), and ABT-199/venetoclax (BCL-2 inhibitor). Data is representative of three more independent experiments

| Cell line | FAB | Age (y), Sex | Mutations | Reference | A-1331852 | S63845 | ABT-263 | ABT-199 |
| --- | --- | --- | --- | --- | --- | --- | --- | --- |
| HL60 | M2 | 36, F | TP53, CDKN2A, NRAS | PMID 2721272 | 12790 | 172.2 | 5724 | 63.37 |
| ML2 | M4 | 26, M | KMT2A-AFDN fusion, NRAS, AFDN-HGNC fusion | PMID 6091813 | 19099 | 299.8 | 4873 | 1636 |
| OCI AML3 | M4 | 57, M | DNMT3A, NRAS, NPM1 | PMID 2538684 and 16079892 | 103464 | 424.1 | 14882 | 22480 |
| Molm-13 | M5a | 20, M | KMT2A-MLLT3 fusion, FLT3-ITD | PMID 9305600 | 83143 | 144062 | 9634 | 22358 |
| MV4-11 | M5 | 10, M | KMT2A-AFF1 fusion, FLT3-ITD | PMID 3496132 | 10901 | 16.85 | 457.6 | 235 |
| THP1 | M5 | 1, M | CSNK2A1-DDX39B fusion, KMT2a, NRAS, TP53 | PMID 60970727 | 27532 | 128.3 | 3282 | 2796 |
| U937 | M5 | 37, M | PICALM-MLLT10 fusion, PTPN11, PTEN, TP53, WT1 | PMID 178611 | 22470 | 609.3 | 13441 | 10757 |
| M-O7E | M7 | 6m, F | ANO7-DHDH fusion, ETO2-GLIS2 fusion, NRAS | PMID 3261598 | 27783 | 10635 | 2512 | 14128 |
| CHRF 288-11 | M7 | 1.5, M | NUP98-KDM5 fusion | PMID 2310825 | 53.65 | 26108 | 2847 | 29329 |

### Metabolic CRISPR screen reveals CI loss enhances sensitivity to MCL-1 inhibition

Due to the metabolic heterogeneity of AML, we sought to identify metabolic lesions that would sensitize *KMT2A*-r AML to MCL-1 inhibition. The MV4-11 cell line was chosen as it is sensitive to MCL-1 inhibition and harbor both *KMT2A*-r and FLT3-ITD mutations, which are known to confer poor prognosis (Supplementary Fig. S1). MV4-11 cells were transduced with a human metabolic CRISPR KO library to selectively ablate genes involved in metabolism and treated with doses of AMG 176, S64315 (also known as MIK665), or AZD5991 that induced approximately 25% cell death (Fig. 1A and B) (ref. 48). After 24 h, the MCL-1 inhibitors were removed, and the cells were further cultured in drug-free media. After 14 population doublings, genomic DNA was isolated, sequenced by Illumina sequencing, and analyzed by MAGeCK VISPR. Several genes were identified that, when lost, conferred sensitivity to MCL-1 inhibitors (Fig. 1C). *SFXN4*, which has been identified as a CI assembly factor, was identified as a common sensitizer amongst all MCL-1 inhibitors (**ref. 67**). Furthermore, the genetic loss of CI components *NDFUB5*, *NDFUB6*, and *NDUFS1* sensitized the MV4-11cells to MCL-1 inhibition by AMG 176 and/or S64315. Loss of *COX17*, a terminal component of the ETC, also sensitized the MV4-11 cells to AMG 176. Enrichr analyses revealed that these genes were generally associated with the function or assembly of CI of the ETC (Fig. 1D, E). Therefore, these data indicate that loss of CI function/assembly potentiates the effect of MCL-1 inhibitors to induce cell death in AML cell lines.

**Figure 1.**
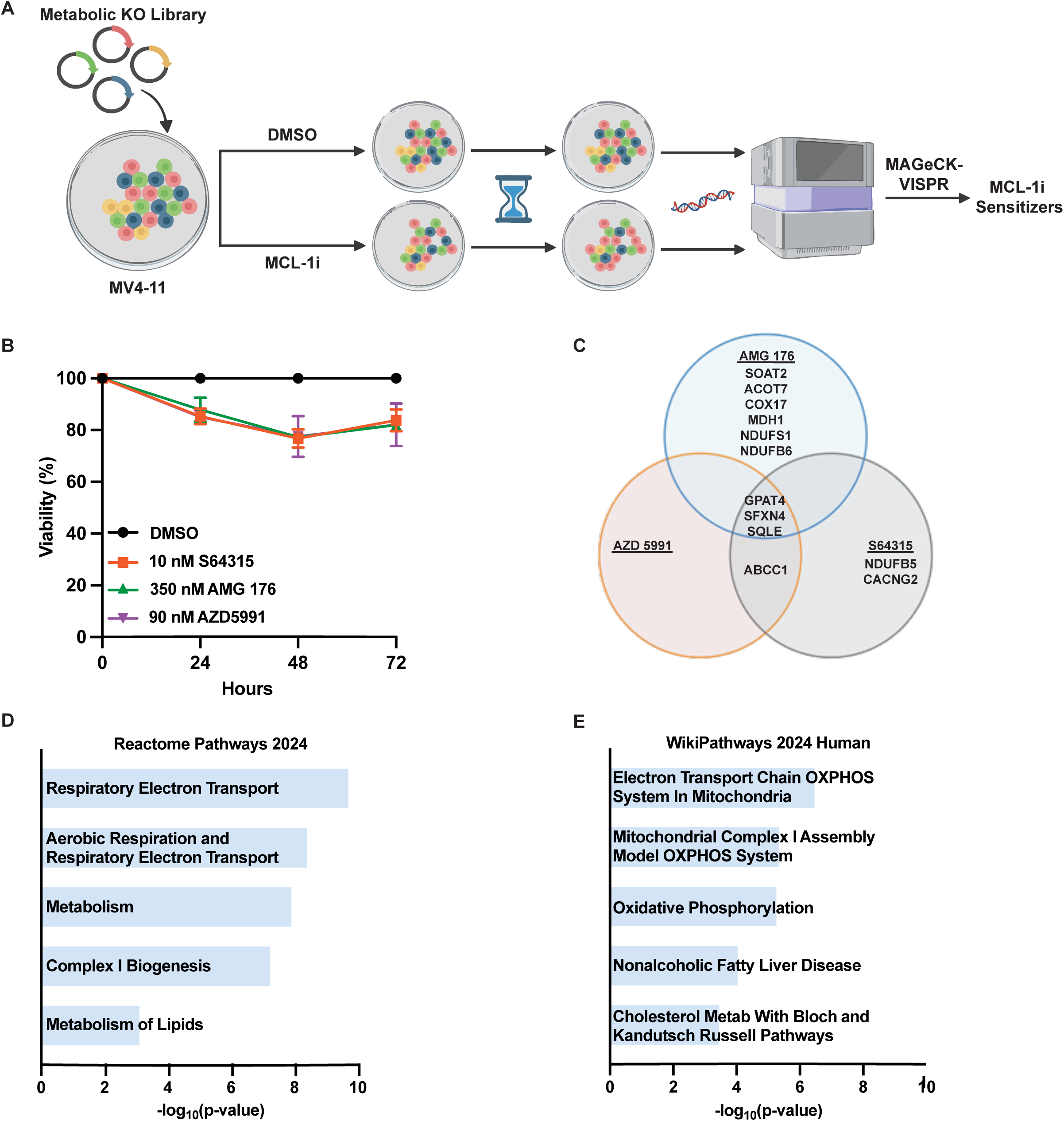
Metabolic CRISPR KO screen identifies OXPHOS as a sensitizer to MCL-1 inhibition in AML. **A,** MV4-11 cells containing the Metabolic CRISPR KO library were treated with either DMSO or low-dose MCL-1i inhibitor for 24 h and then cultured for up to 14 population doublings in drug free media. Illumina sequencing was performed on DNA extracted from the surviving population to determine genes that could be targeted to sensitize AML to MCL-1 inhibition **B,** Cell viability (AV^-^/PI^-^) by flow cytometry of MV4-11 cells treated with indicated doses of AMG 176, AZD5991, or S64315 that resulted in approximately 25% cell death. Data are an average of two or more experiments, and error bars indicate the standard error of the mean (SEM). **C,** MAGeCK-VISPR was used to identify loss of the metabolic related genes that resulted in sensitivity to MCL-1 inhibition. Listed hits from the screen have a false discovery rate of approximately 20% or less at either the early or late timepoint. **D and E,** Enrichr analysis of the pathways that were detected from the combined hits from Fig. 1C.

### MCL-1 inhibitors synergize with the CI Inhibitor, IACS-010759, to induce apoptosis

To pharmacologically validate the CRISPR screen hits, we used the selective CI inhibitor IACS-010759 (hereafter referred to as IACS) that selectively binds to the ND1 subunit of CI and represses OXPHOS (ref. 26, 68). IACS has been tested in clinical trials in both heme and solid cancers as a single agent, exhibiting modest anti-tumor activity at tolerated doses (ref. 25). All three *KMT2A*-r AML cell lines were resistant to IACS treatment-induced cell death, with only minimal cell death induced at the highest doses after 24 h (Fig. 2A). The Seahorse Cell Mito Stress Test was used to measure the amounts of ATP derived from OXPHOS (mitoATP) or from glycolysis (glycoATP). AML cells treated with 50 nM IACS for one hour before the mito stress test had repressed OXPHOS, as shown by decreased production of mitoATP and forced the cells to almost exclusively produce glycoATP (Fig. 2B). These results indicate that IACS can rapidly inhibit OXPHOS without inducing cell death as a single agent.

**Figure 2.**
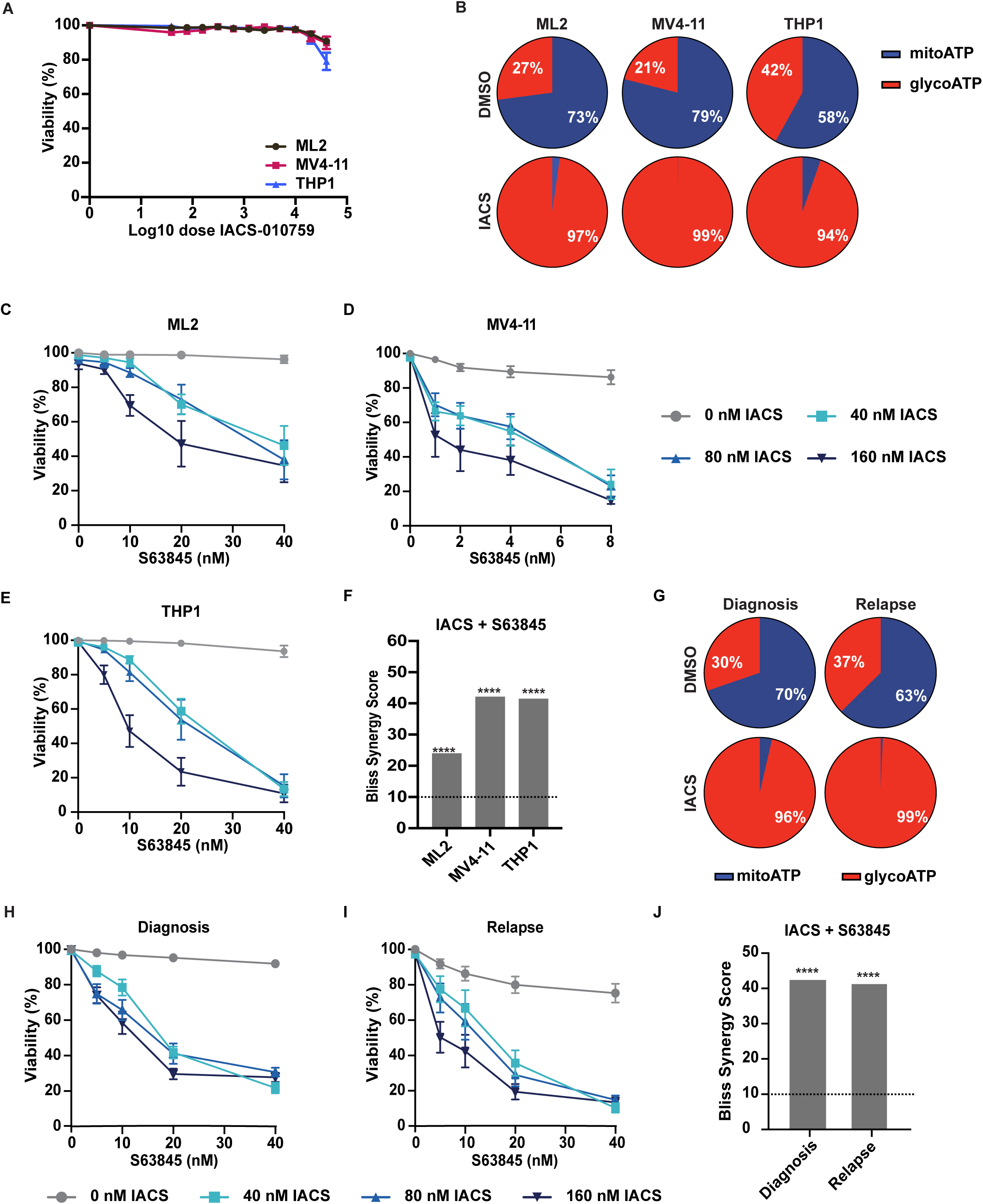
MCL-1 inhibitors synergize with the CI inhibitor, IACS-010759. **A,** Cell viability (AV^-^/PI^-^) of AML cell lines treated with the indicated cells doses of IACS for 24 h. Graphs represent data from three or more independent experiments, and error bars indicate the SEM. **B,** ATP production of ML2, MV4-11, and THP1 cells treated with DMSO or 50 nM IACS for 1 h. **C-E,** Cell viability (AV^-^/PI^-^) of AML cell lines treated with the indicated cells doses of IACS combined with S63845 for 24 h. Graphs represent data from three or more independent experiments and error bars indicate SEM. **F,** Bliss synergy scores derived from the data presented in **C-E**. P-values associated with Bliss synergy scores are calculated through SynergyFinder+ software. P-values < 0.0001 are indicated by ****. Experiments were performed in at least 3 replicates. **G,** ATP production of PDXs derived from two AML patients at disease diagnosis or relapse and treated *ex vivo* with 50 nM IACS or DMSO for 1 h. Data are represented by n=4 mice per group. **H and I,** Cell viability (AV^-^/PI^-^) of PDXs derived from two AML patients at disease diagnosis **(H)** or relapse (**I)** and treated *ex vivo* with the indicated doses of IACS and S63845 for 24 h. Data are represented by n=4 mice per group and error bars indicate the SEM. **J** Bliss synergy scores of the diagnosis or relapse PDX cells treated *ex vivo* with combined doses of S63845 and IACS. Data is representative of n=4 mice. P-values associated with Bliss synergy scores are calculated through SynergyFinder+ software. P-values < 0.0001 are indicated by ****.

To assess whether repression of OXPHOS induced by CI inhibition could potentiate the action of MCL-1 inhibitors, three different *KMT2A*-r AML cell lines were treated with four MCL-1 inhibitors (S63845, AZD5991, AMG 176, or S64315) with or without IACS (Fig. 2C-F and Supplementary Fig. S2A-I). While minimal cell death was observed in cells treated with MCL-1 inhibitors as single agents, the combination treatments significantly reduced cell viability in all cell lines after 24 h of treatment. To assess whether the effect of the combinations were synergistic, the combination drug response was quantified using the Bliss synergy model, where values >10 indicate a synergistic drug interaction (ref. 58). In all three *KMT2A*-r AML cell lines, we observed a significant synergistic interaction when any of the MCL-1 inhibitors were combined with IACS (Fig. 2F and Supplementary Fig. S2J-L).

To expand our analyses beyond AML cell lines, *KMT2A*-r patient-derived xenografts (PDXs) that originated from two patients at either a diagnosis or relapsed disease state were expanded in NSG-SGM3 mice and isolated from the bone marrow for treatment *ex vivo*. Seahorse Cell Mito Stress Test analyses of both *KMT2A*-r PDX samples showed that 1 h of pretreatment with 50 nM of IACS repressed mitoATP production, indicating repression of OXPHOS (Fig. 2G). When IACS and MCL-1 inhibitors were combined, we observed more cell death induced by the combination treatments *ex vivo* as compared to single agents (Fig. 2H and I). Combining IACS with MCL-1 inhibitors resulted in statistically significant synergistic killing by Bliss analysis (**Fig. 2J**). Taken together, these results suggest that repression of CI of the ETC renders *KMT2A*-r AML models more susceptible to cell death by MCL-1 inhibition.

MCL-1 inhibitors trigger intrinsic apoptosis by inducing a BAX- and BAK-dependent mitochondrial outer membrane permeabilization (MOMP) (ref. 69). To test if the cell death observed in response to the combination of IACS and MCL-1 inhibitors was apoptotic, MV4-11 cells lacking both the apoptotic effectors BAK and BAX (hereafter referred to as DKO cells) were generated by CRISPR gene editing (Supplementary Fig. S3A). To assess whether the loss of BAX and BAK compromised the ability of IACS to repress mitochondrial ATP production, WT and DKO MV4-11 cells were pretreated with 50 nM IACS for 1 h before assessing mitochondrial function by Seahorse XF Cell Mito Stress Test. IACS treatment inhibited mitoATP production similarly in both the WT and DKO MV4-11 cells (Supplementary Fig. S3B). Furthermore, when DKO cells were treated with IACS combined with MCL-1 inhibitors, they were resistant to the combination treatment (Supplementary Fig. S3C-F). Bliss synergy analyses of the responses (Supplementary Fig. S3C) failed to show synergy when compared to the WT MV4-11 cells (Supplementary Fig. S2K). The decrease in Bliss synergy scores observed in the DKO cells was shown to be significant as determined by the Zp-value generated using the Z-test between Bliss synergy scores and the corresponding p-value of the WT and DKO cells. These data demonstrate that the pro-apoptotic effectors BAK and BAX are required to mediate cell death in response to the combination treatment, indicating the canonical apoptosis pathway is engaged by the combination therapy.

### IACS primes the mitochondria for apoptosis

Mitochondrial priming describes how near a cell is to reaching the apoptotic threshold; specifically, how “ready” its mitochondria are to undergo MOMP in response to a pro-apoptotic signal (ref. 70). To better understand the mechanism of how IACS potentiates the effects of MCL-1 inhibitors, we tested the effects of IACS on mitochondrial priming using dynamic BH3 (dBH3) profiling, which tests drug-induced MOMP through the observation of Cytochrome *c* (Cyt *c*) release (ref. 71). Three *KMT2A*-r AML cell lines were treated with IACS for 24 h, permeabilized with digitonin, and then exposed to the BIM peptide. The BIM peptide mimics its BH3 domain that binds to all the anti-apoptotic BCL-2 family members and causes Cyt *c* release (ref. 72). Titration of the BIM peptide up to 10 μM revealed that Cyt *c* was more readily released from the mitochondria after IACS treatment when compared to cells treated with DMSO (Fig. 3A-C). The readiness of IACS-treated samples to release Cyt *c* indicates that the cells were more primed after culture in IACS.

**Figure 3.**
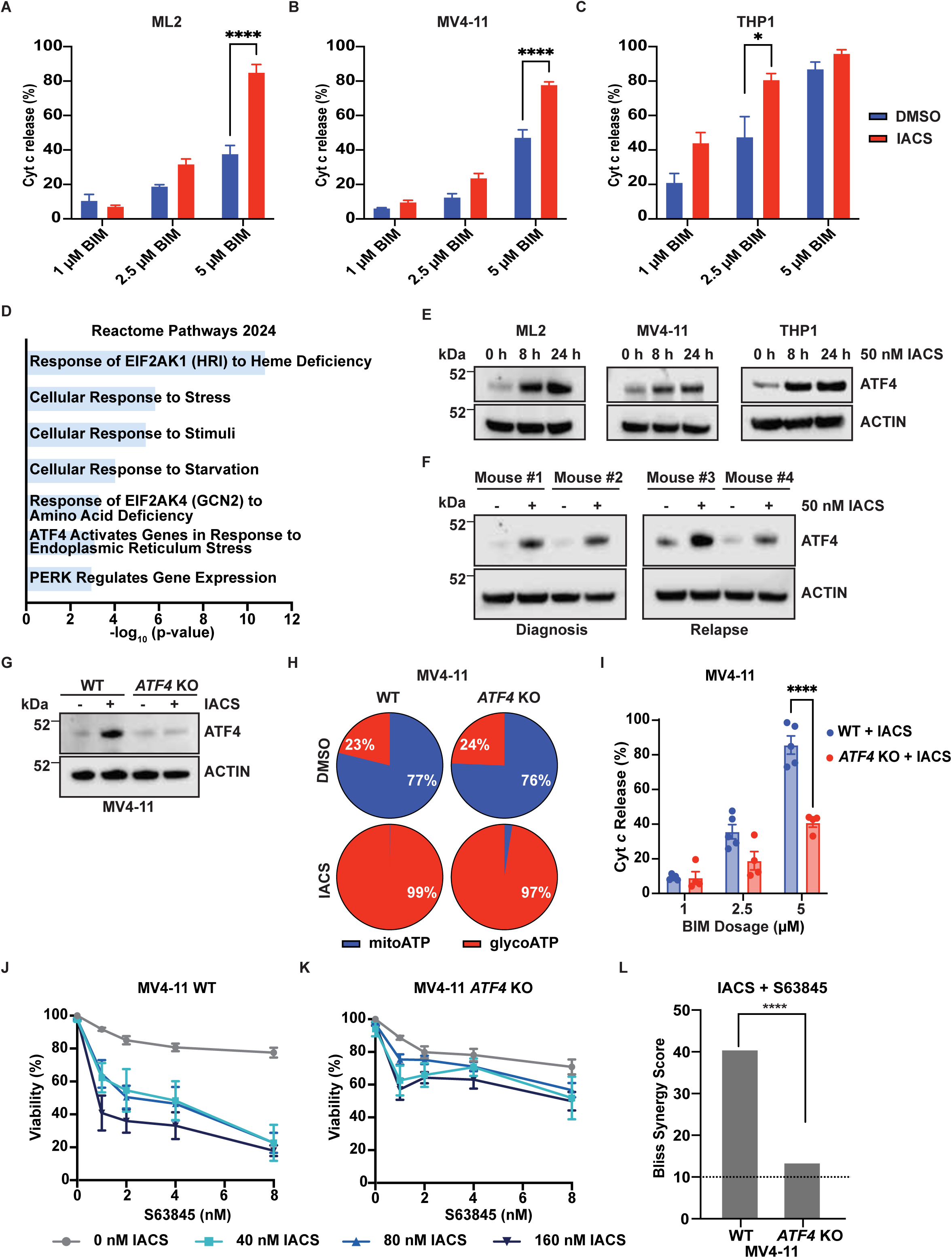
IACS + MCL-1i induces cell death through ATF4 activation. **A-C,** Dynamic BH3 profiling of **(A)** ML2, **(B)** MV4-11, and **(C)** THP1 cells treated with 50 nM IACS for 24 h. Statistical analysis performed using 2way ANOVA of n=3 experiments, and error bars indicate the SEM. P-values < 0.05 are indicated by * and p-values ≤ 0.0001 indicated are by ****. **D,** Enrichr analysis of the top 30 upregulated indexed hits identified by RNA sequencing after IACS treatment in MV4-11 and THP1 cells. **E,** Western blot analysis of ATF4 activation in AML cell lines treated with 50 nM IACS for 8 or 24 h. **F,** Western blot analysis of PDX cells treated with 50 nM IACS for 24 h *ex vivo* **G,** Western blot of WT and *ATF4* KO MV4-11 cell lines treated with IACS for 24 h. **H,** ATP production graphs of WT and *ATF4* KO MV4-11 cells treated with IACS for 1 h. **I,** dBH3 profiling of MV4-11 WT or *ATF4* KO treated cells with IACS. Error bars are represented as the SEM. Two-way ANOVA statistical analysis performed on n=3 or more dBH3 profiling experiments. Cell viability graphs (AV^-^/PI^-^) of WT **(J)** and *ATF4* KO (**K)** MV4-11 cells treated with increasing doses of IACS and S63845. Graphs represent data from three or more independent experiments, and error bars indicate the SEM. **L,** Bliss synergy scores of WT and *ATF4* KO. Dashed line indicates values over 10 as synergistic. Statistical analysis shows Zp-value comparing Bliss scores and p-values between the WT and *ATF4* KO. Zp-values ≤ 0.0001 are indicated by ****.

### IACS induces the integrated stress response (ISR) to activate ATF4

To mechanistically interrogate how IACS primes *KMT2A*-r AML cells to the effects of MCL-1 inhibition, THP1 and MV4-11 cell lines were treated with 50 nM IACS for either 8 or 24 h and RNA sequencing was performed. Enrichr analysis of the top 30 indexed hits revealed that IACS treatment triggered transcript expression patterns associated with the ISR; including those associated with the HRI (gene name *EIF2AK1*), GCN2 (gene name *EIF2AK4*), and PERK (gene name *EIF2AK3*) kinase activated pathways (Fig. 3D). Additionally, many of the top indexed hits were either predicted or experimentally validated ATF4 target genes (Table S1). Taken together, the RNA sequencing indicated that IACS activates a genetic pathway downstream of ATF4. To verify the activation of the ISR after IACS treatment, the *KMT2A*-r AML cell lines were treated with 50 nM IACS for 8 or 24 h and western blotted to assess ATF4 induction. ATF4 expression was increased in all three AML cell lines after IACS treatment (Fig. 3E). Furthermore, ATF4 was also induced in the *KMT2A*-r PDX cells treated with IACS *ex vivo* for 24 h (Fig. 3F). These data indicate that IACS treatment induces ATF4 activation and the ISR.

To investigate if activation of ATF4 by the co-treatment of IACS and MCL-1 inhibitors had a role in apoptosis, *ATF4* KO MV4-11 cells were generated via CRISPR gene editing (Fig. 3G). To ensure the loss of ATF4 still rendered the cells responsive to IACS, MV4-11 WT and *ATF4* KO cells were treated with 50 nM IACS for 1 h, and cellular respiration was assessed. Even in the absence of ATF4, IACS treatment decreased mitoATP production (Fig. 3H), indicating that IACS inhibits CI independently of the induction of an ATF4-mediated response. To determine whether ATF4 contributed to IACS-induced MOMP, dBH3 profiling was performed to compare the response to IACS in WT and *ATF4* null cells. The absence of ATF4 decreased the Cyt *c* release compared to the WT after IACS exposure (Fig. 3I), indicating a reduction in overall mitochondrial priming. Since the genetic ablation of *ATF4* made the cells less primed for apoptosis after IACS treatment, we hypothesized that the loss of ATF4 would promote resistance to the co-treatment of IACS and MCL-1 inhibitors. The loss of *ATF4* rendered MV4-11 cells less responsive than the WT to the combination treatment of IACS and MCL-1 inhibitor (Fig. 3J and K) and significantly decreased the Bliss synergy score indicating less synergy (Fig. 3L). These results suggest that the induction of ATF4 triggered by IACS treatment is responsible for promoting an apoptotic priming to MCL-1 inhibitors in *KMT2A*-r AML cells.

### IACS activates ATF4 primarily through the DELE1-HRI pathway

To determine the mechanism by which IACS stimulates mitochondrial priming we sought to identify genes that when lost conferred resistance to the co-treatment of IACS and MCL-1 inhibitors. An ISR-focused CRISPR KO library was generated containing guides targeting the four eIF2α kinases, pro-apoptotic genes, and the top 30 upregulated and 5 downregulated transcripts identified by the RNA sequencing, which included 25 predicted or validated ATF4 target genes (Table S1). The ISR KO library was transduced into MV4-11 cells and treated with a combination of 160 nM IACS and 4 nM S64315, which resulted in ∼70% cell death (Fig. 4A). The drugs were washed out after 24 h and cultured in drug-free media for 13 population doublings. MAGeCK VISPR analysis revealed that the loss of the eIF2α kinase, *EIF2AK1* (encodes heme-regulated inhibitor, HRI), *DELE1* (HRI activator), *HSPA5* (ER chaperone), and *ALAS1* (heme biosynthesis enzyme) were significant genes that when lost promoted drug resistance (Fig. 4B). Together, the ISR CRISPR screen implicated the DELE-HRI axis of the ISR as key pro-death signaling pathway when IACS and MCL-1 inhibitors are combined.

**Figure 4.**
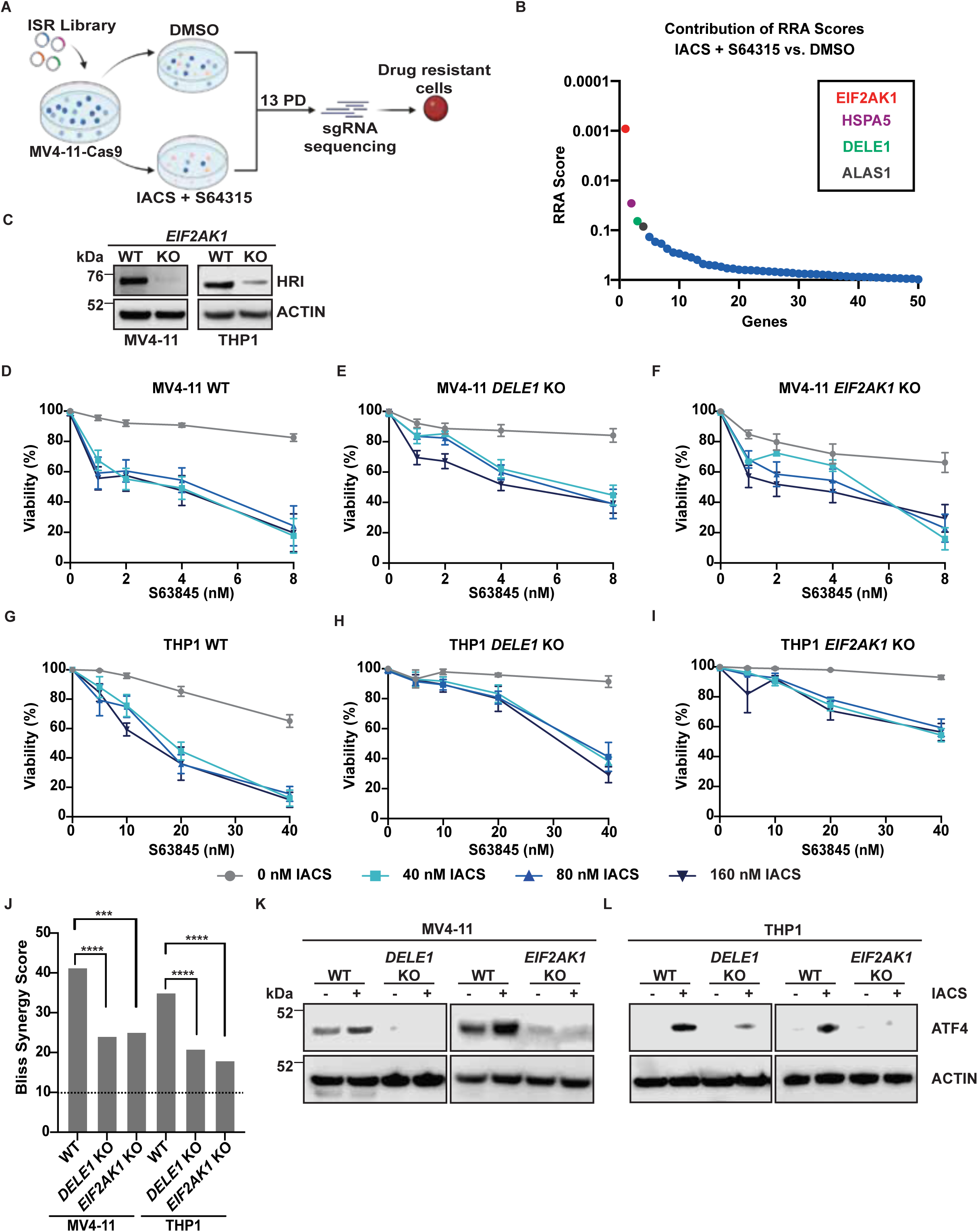
ISR focused CRISPR KO screen identifies loss of DELE1-HRI axis promotes combination therapy resistance. **A,** Schematic of screen showing Cas9 expressing MV4-11 cells transduced with the ISR CRISPR KO library were treated with either DMSO or co-treated with 4 nM S64315 and 160 nM IACS. **B,** Contribution of RRA scores in the combined treatment of S64315 and IACS vs. DMSO treated samples. **C,** Expression levels of HRI in MV4-11 and THP1 KO cell lines. Cell viability (AV^-^/PI^-^) graphs of WT **(D and G)**, *DELE1* KO **(E and H)**, and *EIF2AK1* KO (**F and I)** MV4-11 **(D-F)** and THP1 **(G-I)** cell lines treated with IACS and S63845. Graphs represent data from three more independent experiments, and error bars indicate the SEM. **J,** Bliss synergy scores corresponding to graphs **D-I**. The dashed line indicates values over 10 as synergistic. Statistical analysis is showing Zp-value comparing Bliss scores and p-values between the WT and KOs. Zp-values < 0.0001 are indicated by ****. **K and L,** ATF4 expression levels after IACS treatment in *DELE1* or *EIF2AK1* KO MV4-11 **(K)** and THP1 **(L)** cells.

DELE1 relays mitochondrial stress to the cytosol by activating the kinase HRI, which initiates the ISR to restore cellular homeostasis (ref. 37, 38). Based on the results from our ISR CRISPR screen, we hypothesized that the DELE1-HRI pathway was responsible for the pro-apoptotic signaling when IACS is combined with MCL-1 inhibitors. To test this possibility, we knocked out *DELE1* or *EIF2AK1* in the MV4-11 and THP1 cell lines using CRISPR gene editing and examined 2-3 independent clones per genotype (Fig. 4C). Since there are no good antibodies to detect DELE1, functional analysis by carbonyl cyanide 4-(trifluoromethoxy) phenylhydrazone (FCCP) treatment in the *KMT2A*-r AML cell lines lacking *DELE1* or *EIF2AK1* failed to induce ATF4 expression (Supplementary Fig. S4A and B), indicating an impaired DELE1-HRI signaling axis. IACS pretreatment for 1 h in either *DELE1* or *EIF2AK1* KO cells resulted in rapid repression of mitoATP production (Supplementary Fig. S5A and B). Like the response observed in the *ATF4* KO cells, this supports the ability of IACS to repress CI, regardless of the downstream activation of the ISR.

Furthermore, the loss of *DELE1* (Fig. 4E and H and Supplementary Fig. 6A,D,E) or *EIF2AK1* (Fig. 4F, I and Supplementary Fig. S6B, F, G) substantially blunted the cell death induced in *KMT2A*-r AML cell lines induced by the combination of IACS and S63845 when compared to the WT cell lines (Fig. 4D and G). Zp-vales calculated from the Bliss synergy scores in both MV4-11 and THP1 cells show the loss of *DELE1* or *EIF2AK*1 (Fig. 4J and Supplementary Fig. S6C and H) caused a significant reduction in the Bliss synergy scores compared to WT (Fig. 4J). Furthermore, the absence of *DELE1* or *EIF2AK1* prevented or blunted the ATF4 activation in response to IACS (Fig. 4K and L and Supplementary Fig. S6I, J). Together, these data suggest the DELE1-HRI-ATF4 signaling pathway induces a pro-apoptotic response when IACS is combined with S63845.

### Co-treatment with IACS and S64315 induces an anti-leukemic response *in vivo*

To explore the translational potential of combining MCL-1 inhibitors with IACS *in vivo*, *KMT2A-MLLT3* AML PDX cells, derived from a patient with relapsed disease, were expanded in NSG-SGM3 mice. Once the mice showed approximately 0.5% human CD45^+^ (hCD45^+^) cells in the peripheral blood, they were randomized for treatment with vehicles, IACS (5 mg/kg by oral gavage and vehicle for S64315, once daily), S64315 (6.5 mg/kg by intraperitoneal injection and vehicle for IACS, once daily), or by combination treatment for a 5-day on/2-day off regimen for two weeks (Fig. 5A). At the end of the second treatment cycle, 3 mice per group were euthanized for analysis. From this cohort, flow cytometric assessment of the bone marrow (BM) showed a significant decrease in the hCD45^+^ cells present in the BM in both the S64315 monotherapy and combination-treated group (Fig. 5B).

**Figure 5.**
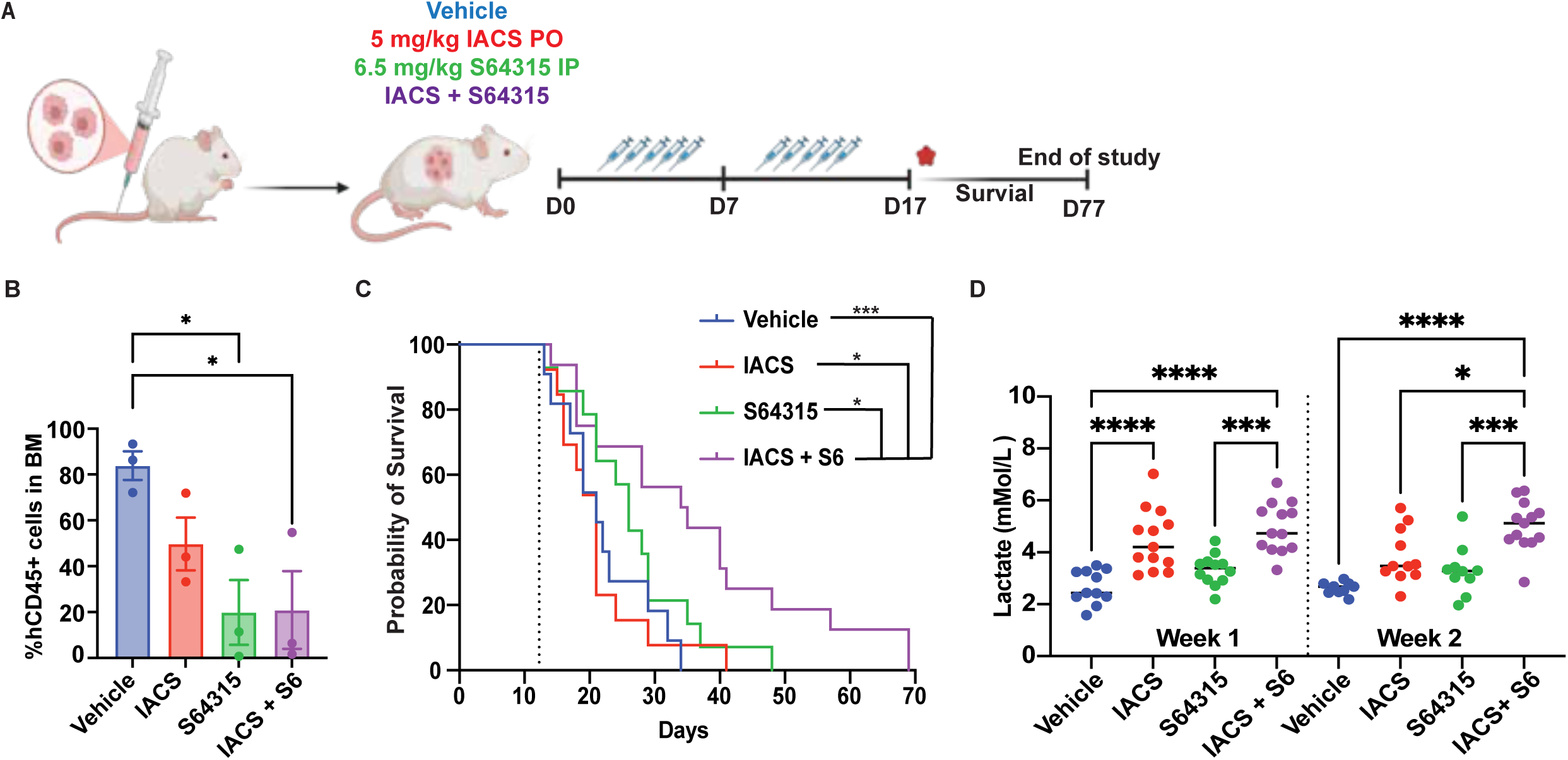
IACS and S64315 induce an anti-leukemic effect *in vivo.* **A,** Schematic of *in vivo* experiments. PDX cells were injected into NSG-SGM3 mice. After engraftment, mice were treated with indicated doses of IACS, S64315, or combination for 5 days on and two days off for two cycles. After the end of the drug cycles, a partial take-down (denoted by red star) was performed of n=3 animals and the remaining animals were kept on the study for survival analyses. **B,** hCD45+ cells in the bone marrow at the end of treatment. n=3 mice per group and error bars are shown as SEM. Statistical significance was determined by ordinary one-way ANOVA. **C,** Survival analysis of mice treated with vehicle, IACS, S64315, or IACS combined with S64315. Statistical significance of the Kaplan-Meier Curve was calculated using the Fleming-Harrington test for right-censored data. The dashed line indicates the last treatment administered (day 12). **D,** Lactate readings from blood plasma samples after one or two weeks on treatment. The line is positioned at the sample median and statistical significance was determined using ordinary one-way ANOVA. P-values < 0.05 are indicated by *, < 0.001 are indicated by ***, <0.0001 are indicated by ****.

After the two treatment cycles, the remaining mice were monitored twice daily for up to two months for survival studies. Once the mice reached a scientific endpoint (i.e. hind leg paresis, tachypneic, delayed response, hunched posture, and weight loss), they were euthanized, and the survival post-treatment was recorded and analyzed. The lifespan of the mice that received the combination therapy was extended significantly with a median survival of 34.5 days (Fig. 5C). The vehicle and IACS cohorts had a median survival of 21 days, indicating a lack of efficacy for IACS as a single agent. Mice that received S64315 treatment had a minor survival advantage over the vehicle and IACS treated cohorts with a median survivals 26 days. Despite the extended survival of the combination-treated mice, all mice eventually exhibited physical symptoms of leukemia and reached the defined scientific endpoint.

IACS use *in vivo* has been reported to induce toxicities during clinical trials including lactic acidosis and peripheral neuropathies (ref. 25). To determine whether our treatment with IACS and/or MCL-1 inhibitors caused any such toxicities *in vivo*, we assessed blood serum lactate levels in at least nine mice per group after one or two weeks on treatment (Fig. 5D). The S64315-treated cohort had similar levels of lactate when compared to DMSO. Elevated lactate levels were observed in both the IACS- and IACS and S64315-treated cohorts, indicating IACS repressed OXPHOS *in vivo.* Mice treated with IACS as either a monotherapy or combination did not reach the 10 mM serum lactate levels seen in metformin-induced models of lactic acidosis (ref. 73). However, mice treated with 5 mg/kg IACS exhibited serum lactate levels comparable to those previously reported, in which peripheral neuropathy was observed (ref. 25).

## Discussion

MCL-1 inhibiting BH3-mimetics constitute a compelling class of anticancer agents because MCL-1 antagonizes apoptosis and drives both tumor maintenance and resistance to conventional therapies (ref. 10, 74, 75, 76, 77). However, the clinical development of MCL-1 inhibitors has been hindered by dose-limiting toxicities that restrict effective drug exposure in phase I trials. To maximize therapeutic benefit while minimizing toxicity, we sought to identify combination strategies that enhance the efficacy of MCL-1 inhibition at lower doses. To identify strategies that potentiate the effects of MCL-1 inhibition, a metabolic CRISPR knockout screen revealed that loss of CI components or assembly factors sensitized AML cells to MCL-1 inhibition. As a proof-of-concept, pharmacologic CI inhibition with IACS synergized with multiple MCL-1 inhibitors to trigger robust apoptosis in *KMT2A-r* AML cell lines and patient-derived xenografts. Mechanistically, IACS increased mitochondrial apoptotic priming and activated the ISR via ATF4, whose loss diminished combination-induced apoptosis. CRISPR screening and genetic validation identified the DELE1-HRI axis as the upstream mediator of IACS-induced ATF4 activation and cell death potentiation. Finally, co-treatment with IACS and an MCL-1 inhibitor extended survival and reduced leukemic burden in *KMT2A-r* AML xenograft models, establishing mitochondrial CI inhibition as a metabolic vulnerability that enhances MCL-1-targeted therapy.

Our findings further demonstrate that the increased apoptotic priming induced by CI inhibition depends on the stress-responsive transcription factor ATF4. Notably, ATF4 plays a dual role in leukemogenesis, acting both as a mediator of leukemic adaptation and as a potential therapeutic effector. Under basal or oncogenic stress conditions, ATF4 supports leukemic cell survival by maintaining amino acid metabolism, redox balance, and mitochondrial homeostasis, thereby promoting leukemogenesis, disease persistence, and therapeutic resistance (ref. 43). In contrast, when excessively or acutely activated by mitochondrial or metabolic stress, ATF4 drives pro-apoptotic signaling that enhances therapeutic responses, particularly in contexts where the ISR is pharmacologically engaged to overcome leukemic resistance (ref. 55, 78, 79). Although ATF4 is known to regulate diverse adaptive responses to metabolic and proteotoxic stress, the downstream mechanisms by which it modulates mitochondrial priming in response to CI inhibition remain unclear. Our RNA sequencing data did not reveal major changes in the transcriptome of the BCL-2 family proteins induced by IACS treatment. Given that post-translational modifications of BCL-2 family proteins can fine-tune the balance between pro- and anti-apoptotic interactions (ref. 80, 81, 82), such modifications may promote alterations in apoptotic priming (ref. 83). Alternatively, an as-yet-unidentified regulator of apoptotic priming may contribute to the modulation of mitochondrial sensitivity following CI inhibition. Together, these findings suggest that ATF4-dependent reprogramming of mitochondrial priming after IACS treatment may involve mechanisms beyond changes in BCL-2 family expression. Further studies are needed to clarify how metabolic stress influences apoptotic susceptibility.

ATF4 can be induced through phosphorylation of eIF2α by any of four stress-responsive kinases, PERK (ER stress), GCN2 (amino acid deprivation), HRI (mitochondrial or heme stress), or PKR (viral infection or double-stranded RNA), each activating the ISR under distinct cellular stress (ref. 84, 85, 86, 87, 88, 89, 90). Mechanistically, we identify that repression of CI function induces the activation of the DELE1-HRI mitochondrial stress pathway that induces ATF4. DELE1, initially identified as a DAP3 binding molecule that promoted apoptosis and later as an activator of HRI-mediated stress signaling has emerged as a central mediator of the mitochondrial stress response in both normal and malignant contexts (ref. 37, 38, 91). In normal tissues, DELE1 primarily supports protective adaptation; studies in skeletal muscle and heart demonstrated that while DELE1 is dispensable under basal conditions, its loss exacerbates pathology in models of mitochondrial myopathy (ref. 92, 93). In cancer, however, DELE1 appears to have context-dependent roles: it is elevated in colorectal cancer and linked to resistance to fluorouracil and oxaliplatin (ref. 94). Yet *DELE1* is frequently deleted in -5/del(5q) AML, where its loss confers resistance to mitochondrial stressors (ref. 95). Our findings indicate that combining IACS with MCL-1 inhibition engages mitochondrial stress through DELE1-HRI signaling, paralleling the resistance observed in *DELE1*-deficient -5/del(5q) leukemia and highlighting DELE1’s capacity to facilitate apoptosis under mitochondrial stress. These results suggest that exploiting DELE1-mediated ATF4 activation could enhance cancer cell death and potentially improve responses to both traditional and targeted therapies.

The mitochondrial stressors CCCP and oligomycin contributed to the discovery of the OMA1-DELE1-HRI arm of the ISR (ref. 37, 38). However, inhibition of ETC complexes can also activate ATF4 through the mitochondrial ISR. While we used IACS to induce ATF4 activation, other CI inhibitors, such as metformin and rotenone, have also been shown to activate ATF4 (ref. 79, 96, 97). Furthermore, inhibition of complex II and complex III similarly trigger ATF4 activation (ref. 98, 99, 100). Although less extensively studied, perturbation of complex IV activity, by agents such as carbon monoxide or arsenic trioxide, can also induce ATF4. Several ETC inhibitors, particularly those targeting CI (metformin, phenformin, IACS, IM156) or complex III (atovaquone) have been tested as cancer therapies, either alone or in combination with standard chemotherapy (ref. 32, 33, 34, 35, 36, 101, 102). Direct inhibition of complex IV remains challenging due to toxicity, though agents that indirectly disrupt mitochondrial function, such as arsenic trioxide, are FDA-approved for certain leukemias. Although we do not explore this question here, we hypothesize that any ETC impairment sufficient to activate ATF4 would synergize with BH3 mimetics. This opens the possibility of multiple potential methods to repress ETC function to improve responses in poor-prognosis AML.

IACS and MCL-1-targeting BH3 mimetics have shown antitumor efficacy against hematologic malignancies in preclinical models; however, their translation to clinical use has been hindered by dose-limiting toxicities. IACS is associated with dose-limiting hematologic adverse events, including neutropenia, thrombocytopenia, and anemia, and has been linked to neurological complications, such as peripheral neuropathy and cognitive impairment, often attributed to lactic acidosis (ref. 25). Consistent with prior studies, administration of IACS, either as a single agent or in combination with MCL-1 inhibition, resulted in elevated serum lactate levels in treated mice (ref. 25); however, these levels did not reach thresholds defined for lactic acidosis in murine models, and no overt peripheral neuropathy was observed (ref. 103, 104). Gross pathology revealed leukemic infiltration of the brain and spine, but no other morphological abnormalities were detected, and cardiomyocyte integrity remained intact, consistent with the reduced affinity of most MCL-1 inhibitors for murine MCL-1 relative to human MCL-1 (ref. 14, 105, 106). Our findings further indicate that brief, low-dose exposure to IACS in combination with S64315 was sufficient to extend survival in mice with *KMT2A-r* AML, suggesting that prolonged high-level IACS exposure may not be required for efficacy. Importantly, this short-course regimen mitigated toxicities historically associated with MCL-1 inhibitors and IACS, including myelosuppression, lactic acidosis, and peripheral neuropathies (ref. 25). A limitation of this study is that extended dosing schedules were not evaluated, which may further enhance combinatorial efficacy but could also increase the risk of adverse events. Moreover, the higher affinity of MCL-1 inhibitors for human compared to murine MCL-1 complicates toxicity assessment in mouse models (ref. 13, 107).

In summary, our results highlight a therapeutic strategy to enhance the efficacy of MCL-1 inhibitors in AML by co-administering IACS to activate a pro-apoptotic ISR. By leveraging mitochondrial stress-induced apoptotic signaling, this combination approach has the potential to overcome resistance mechanisms and selectively sensitize AML cells to BH3-mimetic therapy. Importantly, this strategy may provide a means to improve treatment outcomes in high-risk or refractory AML, a patient population with limited therapeutic options. Future studies are warranted to validate these findings in additional *in vivo* models, define the underlying molecular mechanisms in greater detail, and assess the safety and translational potential of this combination for clinical application. Such investigations will be critical in establishing establish the feasibility of targeting stress-induced apoptosis as a complementary approach to existing AML therapies.

## Data availability

CRISPR KO screen raw sequencing data is archived in NIH BioProject under Accession SUB15754309 and will be made publicly available upon publication. The RNA sequencing data has been deposited in NCBI’s Gene Expression Omnibus and are accessible through GEO Series accession number GSE309897 and will be made publicly available upon publication.

## Acknowledgements

We thank members of the Opferman lab for their insight and support with this project. We thank Danny D’Amore for editorial assistance. We thank Elizabeth D. Arnold and Morgan Reynolds-Gagliano from the Center for Advanced Genome Engineering (CAGE) for their assistance in the generation of mutant cell lines. We thank Jon P. Connelly from the CAGE for his help with the generation of the ISR library and CRISPR guidance. We thank Paula Perez Sanchez and the rest of the Center of Excellence for Leukemia Study (CELs) Flow Cytometry Core for their assistance with the flow cytometry of PDX models. We would like to thank the Preclinical Therapeutics Program (PTP) for their help with preclinical studies. This research included experiments conducted by the Center for Advanced Genome Engineering, which is supported in part the National Cancer Institute grant P30 CA021765. This research included reagents provided by the St. Jude Vector Laboratory Shared Resource. The preclinical formulation utilized in this publication was provided by the Analytical Technology Center-Preclinical Formulation Team (ATC-PFT). This research is funded by the American Lebanese Syrian Associated Charities.

## Authors’ contributions

**L. Brakefield-Laird:** Conceptualization, data curation, formal analysis, investigation, methodology, project administration, resources, supervision, validation, visualization, writing-original draft, writing-review and editing. **A. Budhraja:** Conceptualization, formal analysis, investigation, methodology, validation, visualization, writing-review and editing. **P. M. Hall:** Data curation, formal analysis, investigation, methodology, resources, writing-review and editing. **D. Brewington:** Investigation, writing-review and editing. **J. Moore:** Investigation, writing-review and editing. **J.Lott:** Formal analysis, investigation, resources, writing-review and editing. **Y. Ni:** Formal analysis, data curation, software, writing-review and editing. **D. Voronin:** Investigation, writing-review and editing. **O. Grant-Chapman:** Investigation, writing-review and editing. **T. Mukiza:** Investigation, writing-review and editing. **T. Wright:** Investigation, writing-review and editing. **Y-D. Wang:** Data curation, formal analysis, software, writing-review and editing. **S. Radko-Juettner:** Conceptualization, resources, writing-review and editing. **S. Pruett-Miller:** Conceptualization, resources, writing-review and editing. **S. Pounds:** Formal analysis, software, writing-review and editing. **P. Vogel** Formal analysis, resources, writing-review and editing. **J.T. Opferman:** Conceptualization, formal analysis, funding acquisition, methodology, project administration, resources, supervision, writing-original draft, writing-review and editing.

## Corresponding author

Correspondence to Joseph T. Opferman

## Ethics Declarations Competing interests

The authors declare that they have no conflicts of interest relevant to this work.

## Ethics approval

Mice used in this study were bred and utilized in compliance with protocols approved by the Institutional Animal Care and Use Committee (IACUC) of SJCRH.

## Supplementary Figures

**Supplementary Figure 1.** *KMT2A*-r AML is sensitive to MCL-1 inhibition. Cell viability (AV^-^ /PI^-^) graphs of a panel of AML cells treated for 24 h with A-1331852, S63845, ABT-263 and ABT-199. Data are an average of three or more experiments and error bars are SEM.

**Supplementary Figure 2.** MCL-1 inhibitors synergize with IACS. Cell viability (AV^-^/PI^-^) of ML2 (**A, D, G)**, MV4-11 **(B, E, H),** and THP1**(C, F, I,)** cells treated with the indicated doses of IACS combined with increasing concentrations of AZD5991 **(A-C)**, AMG 176 **(D-F)**, and S64315 **(G-I)**. Bliss synergy scores of ML2 **(J)**, MV4-11 **(K)**, and THP1 **(L)** treated with IACS and AZD5991, AMG 176, or S64315. Graphs represent data from three or more independent experiments, and error bars indicate the SEM. P-values associated with Bliss synergy scores are calculated through SynergyFinder+ software. P-values < 0.0001 are indicated by ****.

**Supplementary Figure 3.** IACS + MCL-1i induce apoptosis in AML cell lines. **A,** Expression levels of BAK and BAX in MV4-11 *BAK*/*BAX* DKO cells. **B,** ATP production graphs of WT and DKO MV4-11 cells treated with 50 nM IACS for 1 h. Data shown is representative of n=3 experiments. **C,** Bliss synergy scores of DKO cells treated with IACS and MCL-1 inhibitors for 24 h. The dashed line indicates values over 10 as synergistic and values between -10 and 10 are additive. Statistical analysis is showing Zp-value comparing Bliss scores and p-values between the WT (Bliss score shown in Fig. 2 F and Supp Fig. 2 K) and KOs. Zp-values < 0.0001 are indicated by ****. **D-F,** Cell viability (AV^-^/PI^-^) of DKO cells treated with IACS combined with S64315, AMG 176 or AZD5991. Graphs represent data from three or more independent experiments, and error bars indicate the SEM.

**Supplementary Figure 4.** Functional validation of *DELE1* and *EIF2AK1* KO.. **A and B,** All MV4-11 (**A**) and THP1 (**B**) *DELE1* and *EIF2AK1* KO clones treated with 10 μM FCCP for 3 h and evaluated for ATF4 expression. Clone (Cl.1) 1 from each panel serves as the representative clone in the primary figures.

**Supplementary Figure 5.** Loss of *DELE1* or *EIF2AK1* does not alter response to IACS. **A and B,** ATP production graphs of MV4-11**(A)** and THP1 **(B)** WT, *DELE1*, and *EIF2AK1* KO clones treated with DMSO or IACS for 1 h. Clone 1 (Cl.1) depicted in these graphs reflect data shown on the primary clones.

**Supplementary Figure 6.** Loss of *DELE1* and *EIF2AK1* promote resistance to IACS and MCL-1 inhibition in multiple clones. Cell Viability (AV^-^/PI^-^) of MV4-11 *DELE1* Cl. 2 **(A)** and *EIF2AK1* KO Cl. 2 **(B)** clones treated with IACS + S63845 for 24 h. **C,** Bliss synergy scored of the additional *DELE1* and *EIF2AK1* KO clones. Statistical analysis is showing Zp-value comparing Bliss scores and p-values between the WT (Bliss score shown in Figure 4J) and additional KO clones. Zp-values < 0.01 are indicated by ** and < 0.0001 are indicated by ****. THP1 *DELE1* Cl. 2 and 3 **(D and E)** and *EIF2AK1* KO Cl. 2 and 3 **(F and G)** clones treated with IACS + S63845 for 24 h. **H,** Bliss synergy scores of the THP1 *DELE1* and *EIF2AK1* KO clones. The dashed line indicates values over 10 as synergistic. Statistical analysis is showing Zp-value comparing Bliss scores and p-values between the WT (Bliss score shown in Figure 4J) and additional KO clones. Zp-values < 0.0001 indicated by ****. Graphs represent data from three or more independent experiments, and error bars indicate the SEM. **I and J,** Western blot analysis of ATF4 activation in the MV4-11 **(I)** and THP1 **(J)** additional clones treated with IACS.

