## Supplemental Figures for "Mitochondrial Integrated Stress Response Activation Creates a Therapeutic Vulnerability to MCL-1 Inhibition in Acute Myeloid Leukemia"

Figure S1. KMT2A-r AML is sensitive to MCL-1 inhibition.

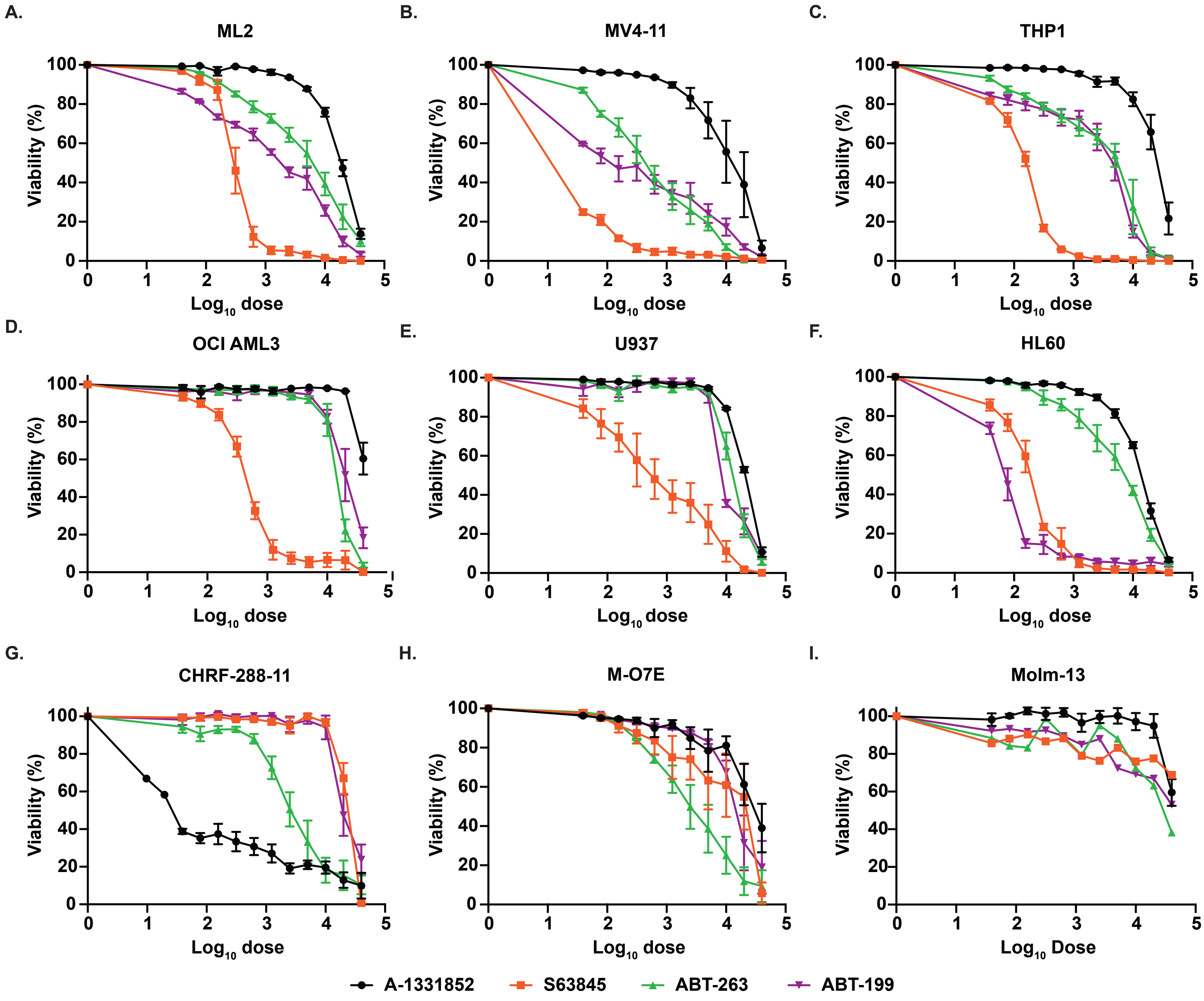

Figure S2. MCL-1 inhibitors Synergize with IACS.

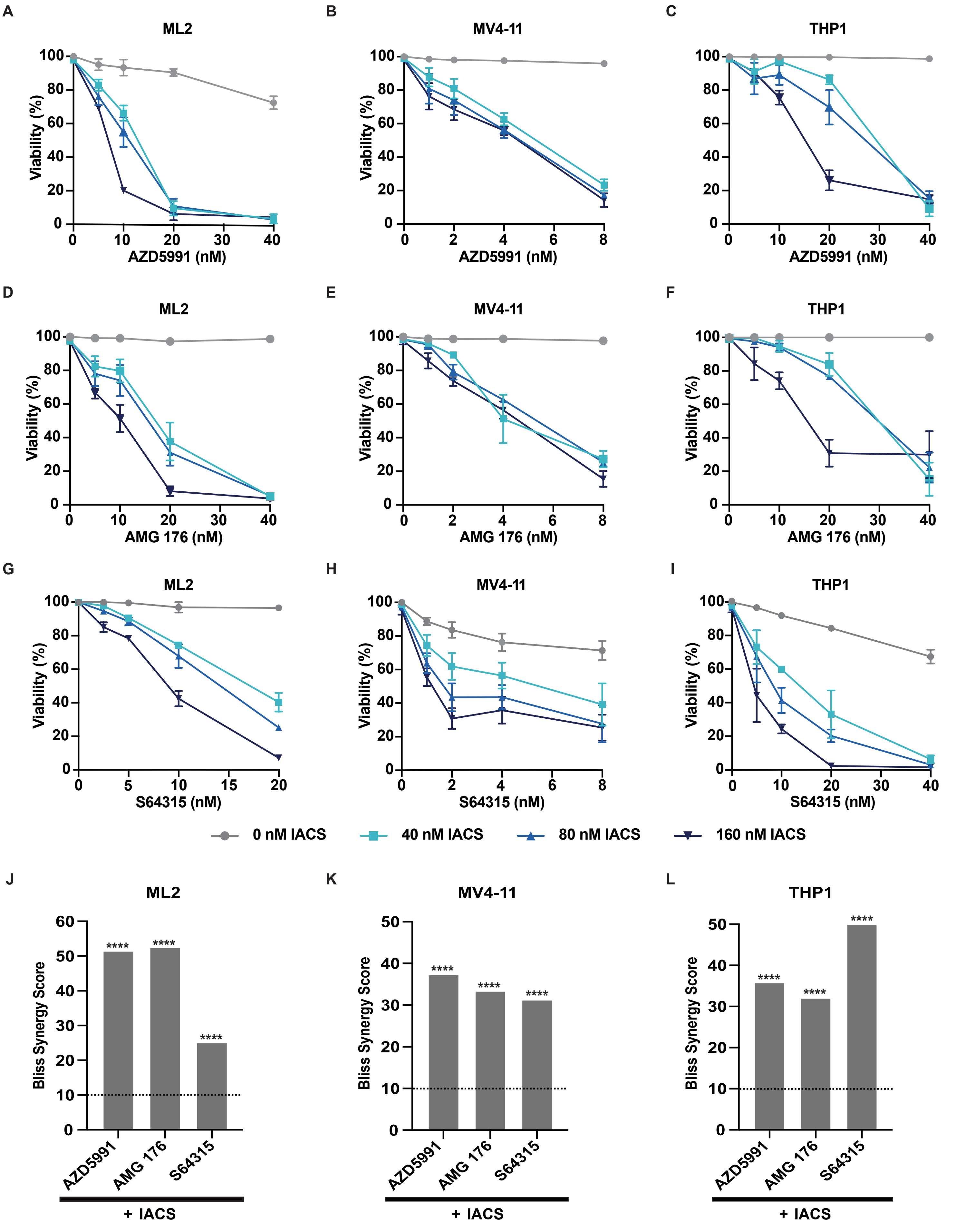

**Figure S3. IACS + MCL-1i induce apoptosis in AML.**

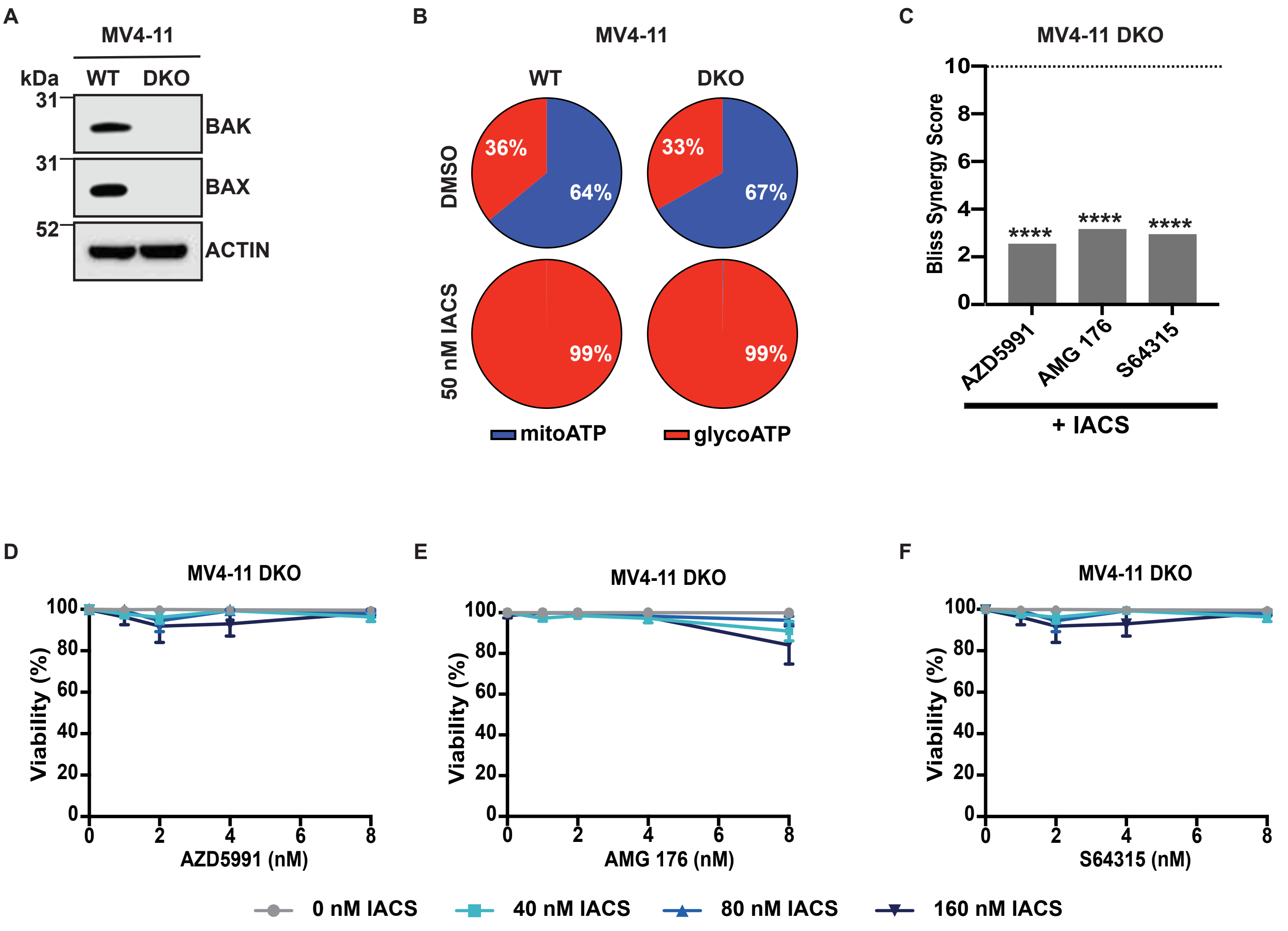

**Figure S4. Functional validation of DELE1 and EIF2AK1 KO.**

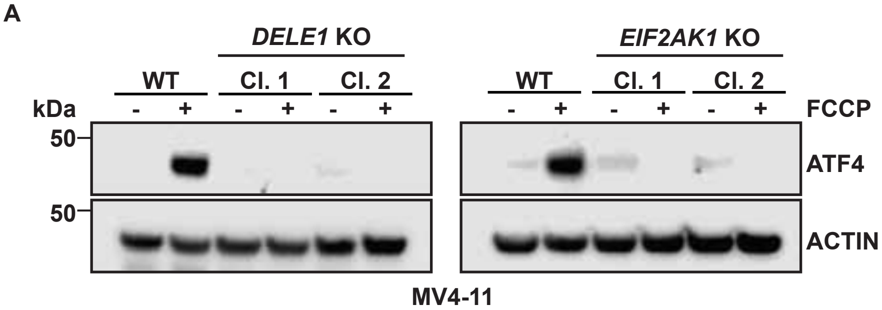


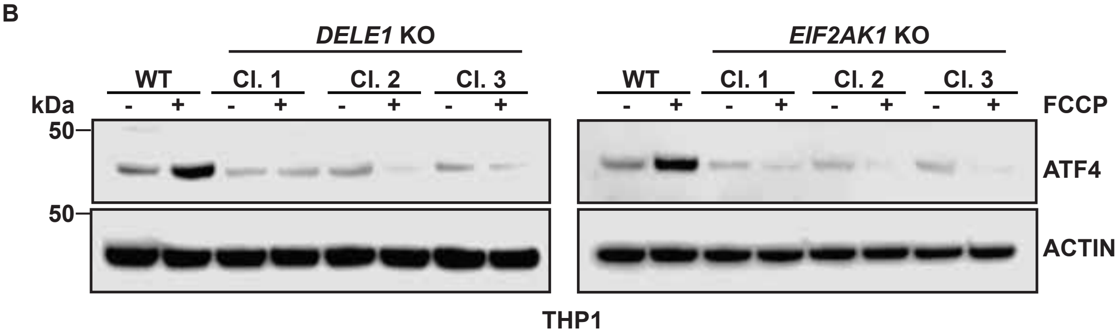

**Figure S5. Loss of DELE1 and EIF2AK1 does not alter response to IACS.**

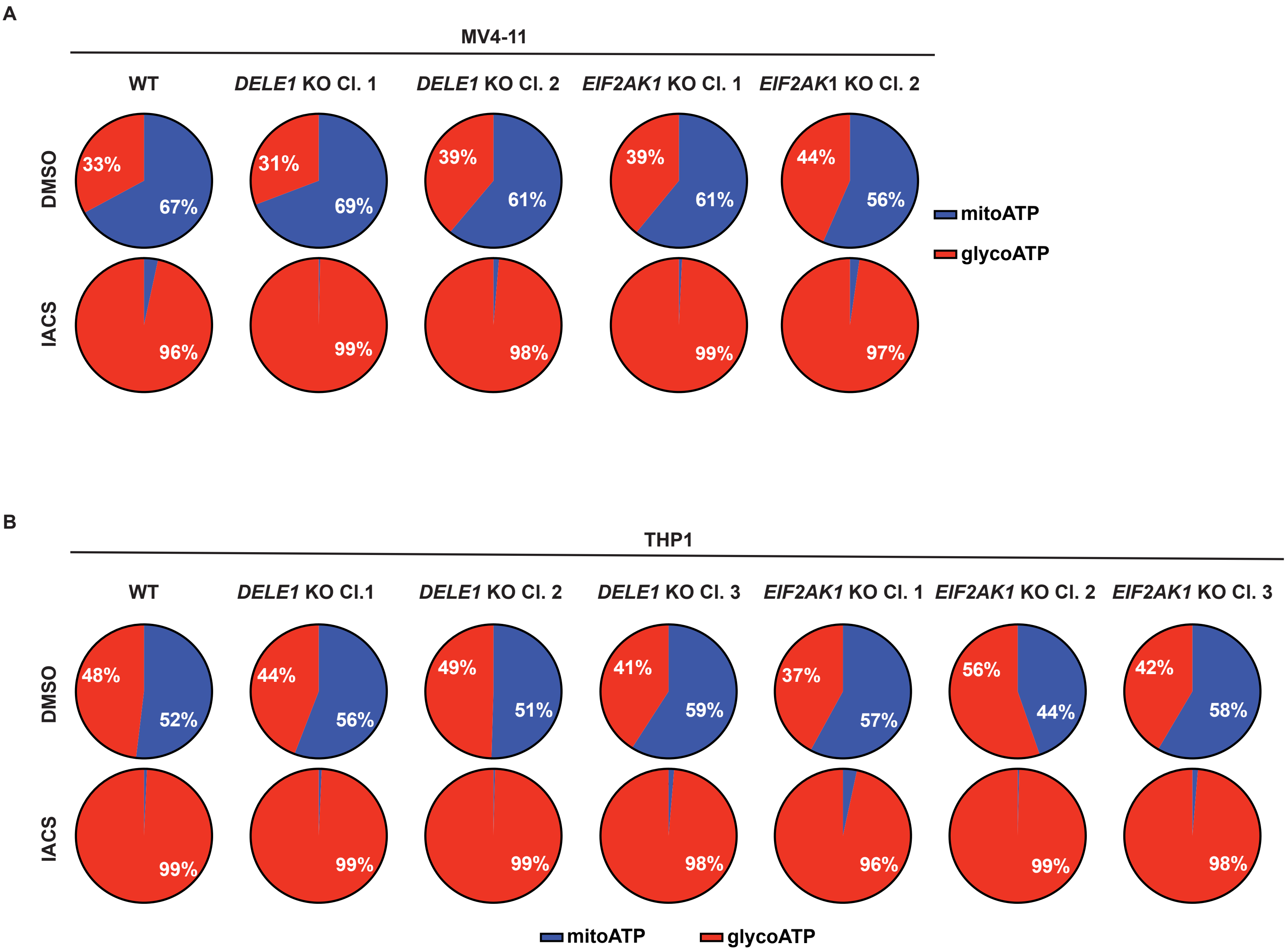

**Figure S6. Loss of DELE1 and EIF2AK1 promote resistance to IACS + MCL-1i in multiple clones.**

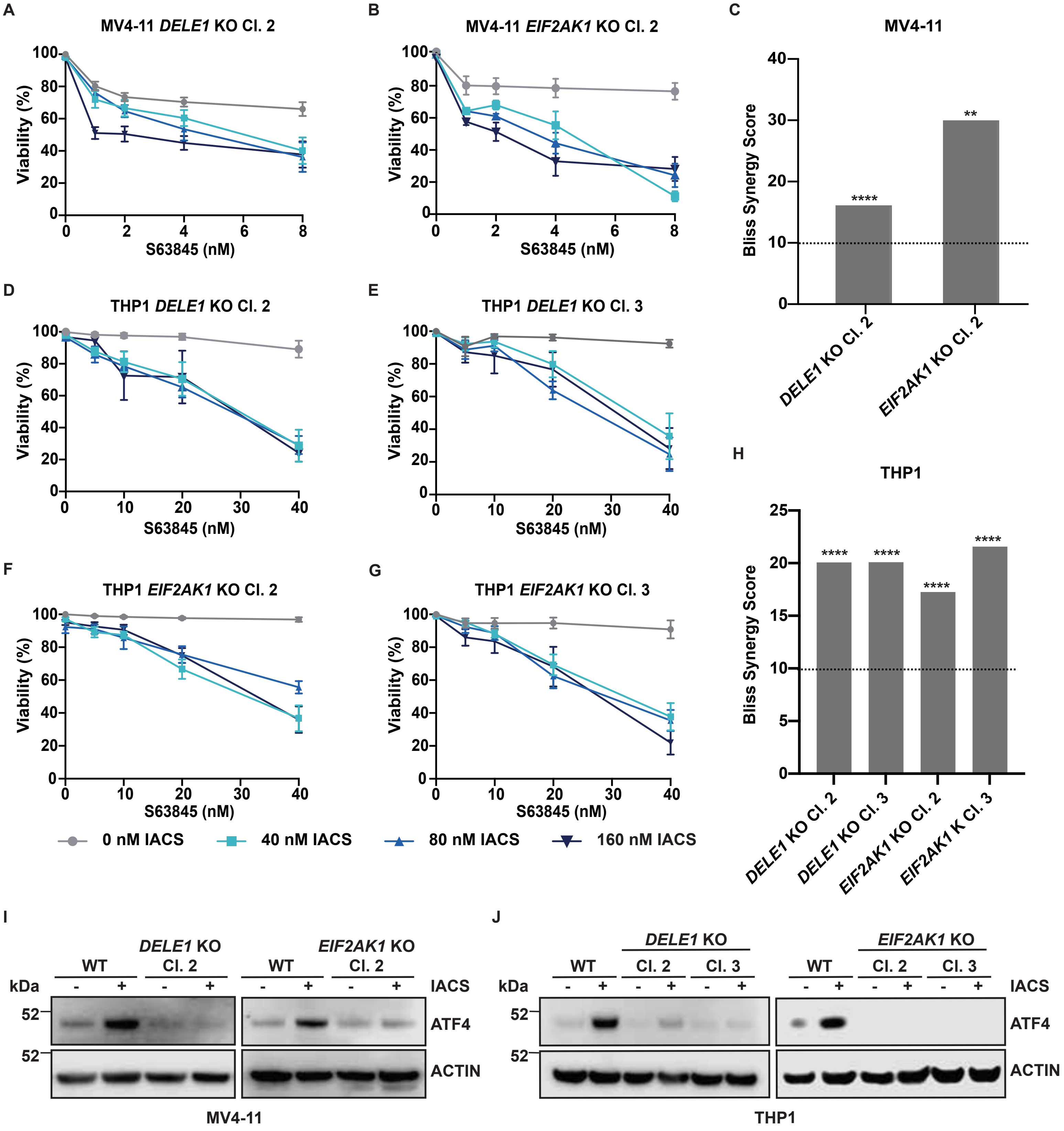
