## Supplementary material for "Mitochondrial Integrated Stress Response Activation Creates a Therapeutic Vulnerability to MCL-1 Inhibition in Acute Myeloid Leukemia": Sup Table 1

**Table S1: ISR KO library guides and sequences.** Guide sequences used to target genes in the ISR CRIPSR KO library.

| **Name** | **gene** | **gRNA sequence** | **Primary pathway** |
| --- | --- | --- | --- |
| **ADM2.g1** | ADM2 | ACCCGCGACCCGTCAAACCCAGG | Predicted or validated ATF4 target |
| **ADM2.g2** | ADM2 | GAGAGGCTGACCCATAACAGGGG | Predicted or validated ATF4 target |
| **ADM2.g3** | ADM2 | GGAGAGGCTGACCCATAACAGGG | Predicted or validated ATF4 target |
| **ADM2.g4** | ADM2 | GGCTGCGGGACAGCGAGCCAGGG | Predicted or validated ATF4 target |
| **ADM2.g5** | ADM2 | ACCGGGCCCTCCAGGCACAGAGG | Predicted or validated ATF4 target |
| **ALAS1.g1** | ALAS1 | GCTGCAGTGGACAATGCCCGAGG | Predicted or validated ATF4 target |
| **ALAS1.g2** | ALAS1 | TTAGTGTGAAAACCGATGGAGGG | Predicted or validated ATF4 target |
| **ALAS1.g3** | ALAS1 | GAAACGATCATACTGAAAAGTGG | Predicted or validated ATF4 target |
| **ALAS1.g4** | ALAS1 | TAACTGCCCCACACACCCGTGGG | Predicted or validated ATF4 target |
| **ALAS1.g5** | ALAS1 | GTCATTGGCCACAAAGCACGAGG | Predicted or validated ATF4 target |
| **ALDH1L2.g1** | ALDH1L2 | ACCCGTTGAAAAGTACTGAGAGG | Predicted or validated ATF4 target |
| **ALDH1L2.g2** | ALDH1L2 | TCTTCTGGCTGGGGTATACGAGG | Predicted or validated ATF4 target |
| **ALDH1L2.g3** | ALDH1L2 | GAACTACCCGCTGATGATGCTGG | Predicted or validated ATF4 target |
| **ALDH1L2.g4** | ALDH1L2 | AACATCCGCCAAAGAAGCGTAGG | Predicted or validated ATF4 target |
| **ALDH1L2.g5** | ALDH1L2 | AAGGTCGTGAGGAAACTGAGAGG | Predicted or validated ATF4 target |
| **APAF1.g1** | APAF1 | TTCCTAAGGAACTCTCCACAGGG | Apoptosis |
| **APAF1.g2** | APAF1 | GTCACCATACATGGAATGGCAGG | Apoptosis |
| **APAF1.g3** | APAF1 | CCAGAGAATACATAACACCTAGG | Apoptosis |
| **APAF1.g4** | APAF1 | AGCATTGTAGAATGATACGTAGG | Apoptosis |
| **APAF1.g5** | APAF1 | ATGGTAAACAGCATCTGTGTGGG | Apoptosis |
| **APBB3.g1** | APBB3 | CAAGTCACTACCTTAGCCCCTGG | APP signaling |
| **APBB3.g2** | APBB3 | TACTACTGGCATGTACCCAGCGG | APP signaling |
| **APBB3.g3** | APBB3 | AGTACGAGGCACTGTATATGGGG | APP signaling |
| **APBB3.g4** | APBB3 | AGTAAGTACCTGCAGCATCGTGG | APP signaling |
| **APBB3.g5** | APBB3 | TCTGGTGCACATCCGTGTGTGGG | APP signaling |
| **ARRDC3.g1** | ARRDC3 | GCCATGTCGGCCTTCGAATGAGG | Arrestin family |
| **ARRDC3.g2** | ARRDC3 | CCTTCGAATGAGGTAGCGAGTGG | Arrestin family |
| **ARRDC3.g3** | ARRDC3 | GCTTGTGGCTAACTTGCGTGGGG | Arrestin family |
| **ARRDC3.g4** | ARRDC3 | GAACTGCTCTTCCCGAATGGTGG | Arrestin family |
| **ARRDC3.g5** | ARRDC3 | CCACTCGCTACCTCATTCGAAGG | Arrestin family |
| **ARRDC4.g1** | ARRDC4 | ATTGGCGACCATGTGTCGAATGG | Arrestin family |
| **ARRDC4.g2** | ARRDC4 | CTTTGGAACAATCAGACGAGAGG | Arrestin family |
| **ARRDC4.g3** | ARRDC4 | GTGTTCGAGGACGAGCGCAAGGG | Arrestin family |
| **ARRDC4.g4** | ARRDC4 | GAAGCATTCAGTACTGTGTGCGG | Arrestin family |
| **ARRDC4.g5** | ARRDC4 | TCCATATTTCCCAGTAAACGAGG | Arrestin family |
| **ASNS.g1** | ASNS | TTGTCATAGAGGGCGTGCAGGGG | Predicted or validated ATF4 target |
| **ASNS.g2** | ASNS | AACGTTTGATGACAGACAGAAGG | Predicted or validated ATF4 target |
| **ASNS.g3** | ASNS | CATTATTAAAAAGGATCCTGAGG | Predicted or validated ATF4 target |
| **ASNS.g4** | ASNS | CTCCATATGTATCTCTACCCAGG | Predicted or validated ATF4 target |
| **ASNS.g5** | ASNS | GTGGTGATCTTCTCTGGAGAAGG | Predicted or validated ATF4 target |
| **BAK1.g1** | BAK1 | ACTTCACCAAGATTGCCACCAGG | Apoptosis |
| **BAK1.g2** | BAK1 | CCTGCTCCTACAGCACCATGGGG | Apoptosis |
| **BAK1.g3** | BAK1 | GCCCACAGCCTGTTTGAGAGTGG | Apoptosis |
| **BAK1.g4** | BAK1 | ACGGCAGCTCGCCATCATCGGGG | Apoptosis |
| **BAK1.g5** | BAK1 | GGTAGACGTGTAGGGCCAGACGG | Apoptosis |
| **BAX.g1** | BAX | TCGGAAAAAGACCTCTCGGGGGG | Apoptosis |
| **BAX.g2** | BAX | GTTTCATCCAGGATCGAGCAGGG | Apoptosis |
| **BAX.g3** | BAX | AGTAGAAAAGGGCGACAACCCGG | Apoptosis |
| **BAX.g4** | BAX | CGAGTGTCTCAAGCGCATCGGGG | Apoptosis |
| **BAX.g5** | BAX | TCTGACGGCAACTTCAACTGGGG | Apoptosis |
| **BBC3.g1** | BBC3 | AGGACACTGCCGAGGGCACCAGG | Apoptosis |
| **BBC3.g2** | BBC3 | GTAGAGGGCCTGGCCCGCGACGG | Apoptosis |
| **BBC3.g3** | BBC3 | TGCTCCGCCAGCGAGAGCGAGGG | Apoptosis |
| **BBC3.g4** | BBC3 | ACCGCCCGCCAGAGCCCCCGGGG | Apoptosis |
| **BBC3.g5** | BBC3 | GGTCCCCGCAGCCGGCCCCGAGG | Apoptosis |
| **CALR.g1** | CALR | TAATCCCCCACTTAGACGGGTGG | Chaperone |
| **CALR.g2** | CALR | ATGAGCAGAACATCGACTGTGGG | Chaperone |
| **CALR.g3** | CALR | TGAGCAGAACATCGACTGTGGGG | Chaperone |
| **CALR.g4** | CALR | GCGGCCAGACAACACCTATGAGG | Chaperone |
| **CALR.g5** | CALR | TCAGGGTTCTGAATCACTGGGGG | Chaperone |
| **CARS1.g1** | CARS1 | GCTTCGATGGACATTCACGGAGG | Predicted or validated ATF4 target |
| **CARS1.g2** | CARS1 | GCCTGAACCACCGAGCCACGTGG | Predicted or validated ATF4 target |
| **CARS1.g3** | CARS1 | CCCTCCAGATGTCTTAACCCGGG | Predicted or validated ATF4 target |
| **CARS1.g4** | CARS1 | ACCCGGGTTAAGACATCTGGAGG | Predicted or validated ATF4 target |
| **CARS1.g5** | CARS1 | ATATTGAAGCGCTGACTCCATGG | Predicted or validated ATF4 target |
| **CHAC1.g1** | CHAC1 | ACGGCGACCCTCAAGCGCTGTGG | Predicted or validated ATF4 target |
| **CHAC1.g2** | CHAC1 | GCTTGGTGGCTACGATACCAAGG | Predicted or validated ATF4 target |
| **CHAC1.g3** | CHAC1 | GCCTACAGCGACAGCCGTGTGGG | Predicted or validated ATF4 target |
| **CHAC1.g4** | CHAC1 | GCCCACACGGCTGTCGCTGTAGG | Predicted or validated ATF4 target |
| **CHAC1.g5** | CHAC1 | GCTCAGTTCCCCCGAAACGACGG | Predicted or validated ATF4 target |
| **CTH.g1** | CTH | AGGCGCCCCTTGCTTGAACGTGG | Predicted or validated ATF4 target |
| **CTH.g2** | CTH | TCTTCAGACCTCGATTGCAGAGG | Predicted or validated ATF4 target |
| **CTH.g3** | CTH | AGCAATTACACCAGAAACCAAGG | Predicted or validated ATF4 target |
| **CTH.g4** | CTH | GCACATATTGTCCATAAGCATGG | Predicted or validated ATF4 target |
| **CTH.g5** | CTH | GCCACGCAGGCGATCCATGTGGG | Predicted or validated ATF4 target |
| **DDIT3.g1** | DDIT3 | CCAGCTGGACAGTGTCCCGAAGG | Predicted or validated ATF4 target |
| **DDIT3.g2** | DDIT3 | TCAGCCAAGCCAGAGAAGCAGGG | Predicted or validated ATF4 target |
| **DDIT3.g3** | DDIT3 | ATTTCCAGGAGGTGAAACATAGG | Predicted or validated ATF4 target |
| **DDIT3.g4** | DDIT3 | CTGGTATGAGGACCTGCAAGAGG | Predicted or validated ATF4 target |
| **DDIT3.g5** | DDIT3 | GACTGGAATCTGGAGAGTGAGGG | Predicted or validated ATF4 target |
| **DDIT4.g1** | DDIT4 | CTGGCCTACAGCGAGCCGTGCGG | Predicted or validated ATF4 target |
| **DDIT4.g2** | DDIT4 | CAGGCCGCACGGCTCGCTGTAGG | Predicted or validated ATF4 target |
| **DDIT4.g3** | DDIT4 | TAGGCATCAGCAGGCGCGCAGGG | Predicted or validated ATF4 target |
| **DDIT4.g4** | DDIT4 | TGAGCGCGGCGGCCGATCTGGGG | Predicted or validated ATF4 target |
| **DDIT4.g5** | DDIT4 | ACAGCGAGCCGTGCGGCCTGCGG | Predicted or validated ATF4 target |
| **DELE1.g1** | DELE1 | TGGGTCTCGTAGTAGGCACCTGG | ISR |
| **DELE1.g2** | DELE1 | TGATAATAAAGGACCGCCTGGGG | ISR |
| **DELE1.g3** | DELE1 | TACCTGTCGAGGTTAGGCACAGG | ISR |
| **DELE1.g4** | DELE1 | GTAGTAGGCACCTGGCATAGCGG | ISR |
| **DELE1.g5** | DELE1 | CAGTAGTAGCATCGAGTCCGAGG | DNase |
| **DNASE2.g1** | DNASE2 | AGCTGGTAGTTATAGACCCAGGG | DNase |
| **DNASE2.g2** | DNASE2 | CTGGTGTTGCTCCGGTACAGCGG | DNase |
| **DNASE2.g3** | DNASE2 | CCGTAGGTACAGGCGCTATGAGG | DNase |
| **DNASE2.g4** | DNASE2 | CAGCTTGTAGACCACGAACCTGG | DNase |
| **DNASE2.g5** | DNASE2 | TCCCCGGACCCTCTAAGAGCTGG | ISR |
| **EIF2AK1.g1** | EIF2AK1 | TGGTGAACTTGAGTCGACCCTGG | ISR |
| **EIF2AK1.g2** | EIF2AK1 | TTGTTGGCTATCACACCGCGTGG | ISR |
| **EIF2AK1.g3** | EIF2AK1 | ATGAACATGTTCTATCCACGCGG | ISR |
| **EIF2AK1.g4** | EIF2AK1 | GGATAGTCGAGAGAAACAAGCGG | ISR |
| **EIF2AK1.g5** | EIF2AK1 | GGCCCGGACCCCGAATATGACGG | ISR |
| **EIF2AK2.g1** | EIF2AK2 | GCAACCTACCTCCTATCATGTGG | ISR |
| **EIF2AK2.g2** | EIF2AK2 | AAGGCCTATGTAATTCCCCATGG | ISR |
| **EIF2AK2.g3** | EIF2AK2 | ATTATGAACAGTGTGCATCGGGG | ISR |
| **EIF2AK2.g4** | EIF2AK2 | CAGGACCTCCACATGATAGGAGG | ISR |
| **EIF2AK2.g5** | EIF2AK2 | AAAGGCAATACGTACCACTGAGG | ISR |
| **EIF2AK3.g1** | EIF2AK3 | TGAAATATCTAACAATGCCCGGG | ISR |
| **EIF2AK3.g2** | EIF2AK3 | TTAGCCAAGCTTGAACACCCGGG | ISR |
| **EIF2AK3.g3** | EIF2AK3 | GAATATACCGAAGTTCAAAGTGG | ISR |
| **EIF2AK3.g4** | EIF2AK3 | CTCAGCGACGCGAGTACCGGCGG | ISR |
| **EIF2AK3.g5** | EIF2AK3 | TATGGACTCAGTGCATATAGTGG | ISR |
| **EIF2AK4.g1** | EIF2AK4 | GCAAGACGACTCCATCGTGGTGG | ISR |
| **EIF2AK4.g2** | EIF2AK4 | AGGATGACCGAGCTGCACGCGGG | ISR |
| **EIF2AK4.g3** | EIF2AK4 | TGTGCACTACCTATACATCCAGG | ISR |
| **EIF2AK4.g4** | EIF2AK4 | GTAGGCCTTCCCATCCACGTTGG | ISR |
| **EIF2AK4.g5** | EIF2AK4 | AACTGGCCAAGAAACACTGTGGG | ISR |
| **GABARAPL1.g1** | GABARAPL1 | GCATTACCAGTAAGGTCAGAGGG | Predicted or validated ATF4 target |
| **GABARAPL1.g2** | GABARAPL1 | AGCATTACCAGTAAGGTCAGAGG | Predicted or validated ATF4 target |
| **GABARAPL1.g3** | GABARAPL1 | CTTGTCCAGATCAGGCACCCTGG | Predicted or validated ATF4 target |
| **GABARAPL1.g4** | GABARAPL1 | GAGCCTTCTCTACAATCACCTGG | Predicted or validated ATF4 target |
| **GABARAPL1.g5** | GABARAPL1 | TCCGGATTAAGAAGTAGAACTGG | Predicted or validated ATF4 target |
| **GDF15.g1** | GDF15 | GGGACGTGACACGACCGCTGCGG | Predicted or validated ATF4 target |
| **GDF15.g2** | GDF15 | AGTTGGTCCGACTGCGACGGCGG | Predicted or validated ATF4 target |
| **GDF15.g3** | GDF15 | GGGCGGCCCGAGAGATACGCAGG | Predicted or validated ATF4 target |
| **GDF15.g4** | GDF15 | AGGGTCCCGGGAAACTTGCGCGG | Predicted or validated ATF4 target |
| **GDF15.g5** | GDF15 | CAGAGCGCGTGCGCGCAACGGGG | Predicted or validated ATF4 target |
| **GRB10.g1** | GRB10 | CCTGGACCTACCTGACAGCGAGG | Predicted or validated ATF4 target |
| **GRB10.g2** | GRB10 | AACATCTTCTCCCTGATCGCTGG | Predicted or validated ATF4 target |
| **GRB10.g3** | GRB10 | CCCTACCTAATCCTAGGTGCGGG | Predicted or validated ATF4 target |
| **GRB10.g4** | GRB10 | CCCGCACCTAGGATTAGGTAGGG | Predicted or validated ATF4 target |
| **GRB10.g5** | GRB10 | CAAGTCGGTCAGACTGTGCGGGG | Predicted or validated ATF4 target |
| **HK3.g1** | HK3 | TAAGGAGACATCGCATTCCGGGG | Predicted or validated ATF4 target |
| **HK3.g2** | HK3 | CTAGGCCTACAACATCGACGTGG | Predicted or validated ATF4 target |
| **HK3.g3** | HK3 | GTCTGCGTCAGCGTCGAGTGGGG | Predicted or validated ATF4 target |
| **HK3.g4** | HK3 | AGCTCAACATCCGAAGCCCCAGG | Predicted or validated ATF4 target |
| **HK3.g5** | HK3 | AGGCAGCATCCGGACCGCAGGGG | Predicted or validated ATF4 target |
| **HSPA5.g1** | HSPA5 | CAGACGGGTCATTCCACGTGCGG | Predicted or validated ATF4 target |
| **HSPA5.g2** | HSPA5 | GGACGGGCTTCATAGTAGACCGG | Predicted or validated ATF4 target |
| **HSPA5.g3** | HSPA5 | CGACATAGGACGGCGTGATGCGG | Predicted or validated ATF4 target |
| **HSPA5.g4** | HSPA5 | GAAGCCCGTCCAGAAAGTGTTGG | Predicted or validated ATF4 target |
| **HSPA5.g5** | HSPA5 | TGTGCAGAAACTCCGGCGCGAGG | Predicted or validated ATF4 target |
| **IL23A.g1** | IL23A | AGGACTCAGGGACAACAGTCAGG | Predicted or validated ATF4 target |
| **IL23A.g2** | IL23A | ACATCCACTAGTGGGACACATGG | Predicted or validated ATF4 target |
| **IL23A.g3** | IL23A | TGCTTGCAAAGGATCCACCAGGG | Predicted or validated ATF4 target |
| **IL23A.g4** | IL23A | ATCCTTTGCAAGCAGAACTAGGG | Predicted or validated ATF4 target |
| **IL23A.g5** | IL23A | GCTGGCACTGAGTCCAGGCAGGG | Predicted or validated ATF4 target |
| **INHBE.g1** | INHBE | GCACAGTTACTGGACAACCGAGG | Predicted or validated ATF4 target |
| **INHBE.g2** | INHBE | CAGTTACTGGACAACCGAGGCGG | Predicted or validated ATF4 target |
| **INHBE.g3** | INHBE | ACCCCAAGCAGAACGAGCTCTGG | Predicted or validated ATF4 target |
| **INHBE.g4** | INHBE | GAGTTAAGGTATGCCAGCCCAGG | Predicted or validated ATF4 target |
| **INHBE.g5** | INHBE | CCCTCCGGAGACTACAGCCAGGG | Predicted or validated ATF4 target |
| **JDP2.g1** | JDP2 | TGTGCCCTCACAGCTAGATGAGG | Predicted or validated ATF4 target |
| **JDP2.g2** | JDP2 | TACGCTGACATCCGCAACCTCGG | Predicted or validated ATF4 target |
| **JDP2.g3** | JDP2 | AAGTCGCAGCAGCCCGATGCCGG | Predicted or validated ATF4 target |
| **JDP2.g4** | JDP2 | AGGGTGCAATCATGGCCCCGAGG | Predicted or validated ATF4 target |
| **JDP2.g5** | JDP2 | TTGGTATACAGGAATCCGAGCGG | Predicted or validated ATF4 target |
| **NDUFA4L2.g1** | NDUFA4L2 | GCAGATCAAAAGACATCCGGGGG | ETC |
| **NDUFA4L2.g2** | NDUFA4L2 | GCCGTGCCTTTACCAGACGTCGG | ETC |
| **NDUFA4L2.g3** | NDUFA4L2 | CGTGCCTTTACCAGACGTCGGGG | ETC |
| **NDUFA4L2.g4** | NDUFA4L2 | GGCAGATCAAAAGACATCCGGGG | ETC |
| **NDUFA4L2.g5** | NDUFA4L2 | GGCTTAATCTGCCTGGGCATGGG | ETC |
| **NTC.g43200** | NTC | AATCGCAGGTATCCCAGAGCNGG | Nontargeting control |
| **NTC.g43242** | NTC | ACGTCAACTGCTGGAGTGGGNGG | Nontargeting control |
| **NTC.g43292** | NTC | AGCTGGACTCTGTAGAAATCNGG | Nontargeting control |
| **NTC.g43336** | NTC | ATACGAGGCGCTTTTCTTTGNGG | Nontargeting control |
| **NTC.g43370** | NTC | ATCTGTCCTAATTCGGATCGNGG | Nontargeting control |
| **NTC.g43458** | NTC | CATTGCACGCCACAGCATTGNGG | Nontargeting control |
| **NTC.g43478** | NTC | CCATCACCGATCGTGAGCCTNGG | Nontargeting control |
| **NTC.g43487** | NTC | CCCGATGGACTATACCGAACNGG | Nontargeting control |
| **NTC.g43535** | NTC | CGACGCTAGGTAACGTAGAGNGG | Nontargeting control |
| **NTC.g43545** | NTC | CGCAATCCCTTAGGATAGCCNGG | Nontargeting control |
| **NTC.g43563** | NTC | CGCGTGCATCTGCCGAAGGCNGG | Nontargeting control |
| **NTC.g43651** | NTC | CTGGTGACCGACAATTACACNGG | Nontargeting control |
| **NTC.g43686** | NTC | GAAGACGTGCTGGCGTCACCNGG | Nontargeting control |
| **NTC.g43715** | NTC | GAGCTTAGCAAAGGGTTGGGNGG | Nontargeting control |
| **NTC.g43812** | NTC | GGAGATGCGGCCTTCTCAAANGG | Nontargeting control |
| **NTC.g43826** | NTC | GGCCGTCGTATTCCCCCAAGNGG | Nontargeting control |
| **NTC.g43840** | NTC | GGGAGTTGATTGTTTCGAGANGG | Nontargeting control |
| **NTC.g43870** | NTC | GGTTAACATCGCCACTCTGANGG | Nontargeting control |
| **NTC.g43914** | NTC | GTGTATGAATGTTAATTCCGNGG | Nontargeting control |
| **NTC.g43968** | NTC | TAGCTCGAGTCATTTCTCTANGG | Nontargeting control |
| **NTC.g44030** | NTC | TCGGGGACCACCCACGATCCNGG | Nontargeting control |
| **NTC.g44081** | NTC | TGGCCACGAATTCCGCCGCCNGG | Nontargeting control |
| **NTC.g44096** | NTC | TGTGCACAAGTCGCAACGAANGG | Nontargeting control |
| **NTC.g44101** | NTC | TTAACTCGAACGCTCGAAAGNGG | Nontargeting control |
| **OMA1.g1** | OMA1 | CTGGAAGTAAGTCCAATCACAGG | ISR |
| **OMA1.g2** | OMA1 | GGAGCAAGCTACTATTATTGGGG | ISR |
| **OMA1.g3** | OMA1 | ACATTAGCATCCACCTCACGGGG | ISR |
| **OMA1.g4** | OMA1 | TTACCAGCAAATGTACTGTATGG | ISR |
| **OMA1.g5** | OMA1 | AAATAGCACATGCAGTACTTGGG | ISR |
| **PCK2.g1** | PCK2 | TGCGTATTATGACCCGACTGGGG | Predicted or validated ATF4 target |
| **PCK2.g2** | PCK2 | GCTGAATGGAAGCACATACATGG | Predicted or validated ATF4 target |
| **PCK2.g3** | PCK2 | CAGCCGAGAGGCGATGCGTAGGG | Predicted or validated ATF4 target |
| **PCK2.g4** | PCK2 | GGCACGAGTAGAGAGCAAGACGG | Predicted or validated ATF4 target |
| **PCK2.g5** | PCK2 | CCAGCCGAGAGGCGATGCGTAGG | Predicted or validated ATF4 target |
| **PHGDH.g1** | PHGDH | TTCATCGAAGCCGTCGCCTGGGG | Predicted or validated ATF4 target |
| **PHGDH.g2** | PHGDH | GTGGTGAACTGTGCCCGTGGAGG | Predicted or validated ATF4 target |
| **PHGDH.g3** | PHGDH | CGTGGAGGGATCGTGGACGAAGG | Predicted or validated ATF4 target |
| **PHGDH.g4** | PHGDH | TGAAGGGGAAATCTCTCACGGGG | Predicted or validated ATF4 target |
| **PHGDH.g5** | PHGDH | ACCAAGGCCCGGTCCCGTGGCGG | Apoptosis |
| **PMAIP1.g1** | PMAIP1 | TCGAGTGTGCTACTCAACTCAGG | Apoptosis |
| **PMAIP1.g2** | PMAIP1 | ACGCTCAACCGAGCCCCGCGCGG | Apoptosis |
| **PMAIP1.g3** | PMAIP1 | CGCTCAACCGAGCCCCGCGCGGG | Apoptosis |
| **PMAIP1.g4** | PMAIP1 | TACCTGCTGGAGCCCGCGCGGGG | Apoptosis |
| **PMAIP1.g5** | PMAIP1 | TTGGAGACAAACTGAACTTCCGG | Apoptosis |
| **RAN.g1** | RAN | ATGGCTATTATATCCAAGGTAGG | Control |
| **RAN.g2** | RAN | GCCGGCCAGGAGAAATTCGGTGG | Control |
| **RAN.g3** | RAN | AACATCCCCATTGTGTTGTGTGG | Control |
| **RAN.g4** | RAN | TGTTGCCACACAACACAATGGGG | Control |
| **RAN.g5** | RAN | AGTCAAATGACGTTTCACGAAGG | Control |
| **RAP1GAP2.g1** | RAP1GAP2 | GTACACAACATTCCGGGACAGGG | GTPase activator |
| **RAP1GAP2.g2** | RAP1GAP2 | ATAGGACACAATCATTTGGGAGG | GTPase activator |
| **RAP1GAP2.g3** | RAP1GAP2 | AACTGAAGACGGTACATGAGCGG | GTPase activator |
| **RAP1GAP2.g4** | RAP1GAP2 | TCCCTTCTAGGACCGGACCAGGG | GTPase activator |
| **RAP1GAP2.g5** | RAP1GAP2 | GAAGCCTCTTACCTTCATGGTGG | ER chaperone |
| **SDF2L1.g1** | SDF2L1 | GGTGACCGGCGTAGAGGCGTCGG | ER chaperone |
| **SDF2L1.g2** | SDF2L1 | GCAATCGGTGACCGGCGTAGAGG | ER chaperone |
| **SDF2L1.g3** | SDF2L1 | AGCGCACTGTCCATAGGTCCAGG | ER chaperone |
| **SDF2L1.g4** | SDF2L1 | ACGACGCCAATAGCTACTGGCGG | ER chaperone |
| **SDF2L1.g5** | SDF2L1 | TCAATACGCACCACCGCGTGCGG | ER chaperone |
| **SESN2.g1** | SESN2 | GATGCCACAGCCAAACACGAAGG | Predicted or validated ATF4 target |
| **SESN2.g2** | SESN2 | CAGGTTGTCTACTCGCCCAGAGG | Predicted or validated ATF4 target |
| **SESN2.g3** | SESN2 | GGACCCGTTGAACAACTCTGGGG | Predicted or validated ATF4 target |
| **SESN2.g4** | SESN2 | GGCCGCCGCTTACCTCTCCGGGG | Predicted or validated ATF4 target |
| **SESN2.g5** | SESN2 | TCGGCCATGGCTCATCACCAAGG | Predicted or validated ATF4 target |
| **SLC7A11.g1** | SLC7A11 | AAGGGCGTGCTCCAGAACACGGG | Predicted or validated ATF4 target |
| **SLC7A11.g2** | SLC7A11 | GAAGAGATTCAAGTATTACGCGG | Predicted or validated ATF4 target |
| **SLC7A11.g3** | SLC7A11 | AGAGGAAAGTCACTTTACTGAGG | Predicted or validated ATF4 target |
| **SLC7A11.g4** | SLC7A11 | TATGCTCTTACCGTATTATGAGG | Predicted or validated ATF4 target |
| **SLC7A11.g5** | SLC7A11 | ATGGATATACATATTGCAAGGGG | Predicted or validated ATF4 target |
| **STC2.g1** | STC2 | CCTCAAGCACGACCTGTGCGCGG | Predicted or validated ATF4 target |
| **STC2.g2** | STC2 | ACACTGAACCTGCACGCTGTGGG | Predicted or validated ATF4 target |
| **STC2.g3** | STC2 | GAACCTGTGCCGCAGAGCGTGGG | Predicted or validated ATF4 target |
| **STC2.g4** | STC2 | AAGTTCACGAGGTCCACGTAGGG | Predicted or validated ATF4 target |
| **STC2.g5** | STC2 | GGAGAACACCCGGGTGATAGTGG | Predicted or validated ATF4 target |
| **TRIB3.g1** | TRIB3 | GACATAGGGCCCAAGACGGGAGG | Predicted or validated ATF4 target |
| **TRIB3.g2** | TRIB3 | CGAGCCACATGCTTGTGCGGGGG | Predicted or validated ATF4 target |
| **TRIB3.g3** | TRIB3 | CCGAGCCACATGCTTGTGCGGGG | Predicted or validated ATF4 target |
| **TRIB3.g4** | TRIB3 | CTGGTGACAGTGCGCCAGGGCGG | Predicted or validated ATF4 target |
| **TRIB3.g5** | TRIB3 | ATCTGGCCCAGTCAGCACGCAGG | Predicted or validated ATF4 target |
| **TSC22D3.g1** | TSC22D3 | AGCGGGGAGAACAACAACCCGGG | Immune/inflammatory regulation |
| **TSC22D3.g2** | TSC22D3 | GCGGCAGGATTCGCTAGAGCCGG | Immune/inflammatory regulation |
| **TSC22D3.g3** | TSC22D3 | TGGACCTGGCCCATCGGAGTGGG | Immune/inflammatory regulation |
| **TSC22D3.g4** | TSC22D3 | GCATGGTCTGGTCGATGTTGCGG | Immune/inflammatory regulation |
| **TSC22D3.g5** | TSC22D3 | CATAGACAACAAGATCGAACAGG | Immune/inflammatory regulation |
| **TXNIP.g1** | TXNIP | GAGATGGTGATCATGAGACCTGG | Predicted or validated ATF4 target |
| **TXNIP.g2** | TXNIP | GTAAGTGTGGCGGGCCACAATGG | Predicted or validated ATF4 target |
| **TXNIP.g3** | TXNIP | CCTGAAAAGGTGTACGGCAGTGG | Predicted or validated ATF4 target |
| **TXNIP.g4** | TXNIP | CCCATCAGGAATGAACATGCAGG | Predicted or validated ATF4 target |
| **TXNIP.g5** | TXNIP | GAAGCGTGTCTTCATAGCGCAGG | Predicted or validated ATF4 target |
| **UTP14C.g1** | UTP14C | ATGTGGACGGACCGAATCCCTGG | Processome |
| **UTP14C.g2** | UTP14C | TAGGGGTTCCGCTTCTTCTGGGG | Processome |
| **UTP14C.g3** | UTP14C | ACAGAAGGTTAATCCAACTGTGG | Processome |
| **UTP14C.g4** | UTP14C | TGATAGGGTCCCATTTGGAGAGG | Processome |
| **UTP14C.g5** | UTP14C | GCATAGCTTGGCGAGCCTCCAGG | Processome |
| **VEGFA.g1** | VEGFA | GGAGGGCAGAATCATCACGAAGG | Predicted or validated ATF4 target |
| **VEGFA.g2** | VEGFA | AGATGTACTCGATCTCATCAGGG | Predicted or validated ATF4 target |
| **VEGFA.g3** | VEGFA | TGGTTTCGGAGGCCCGACCGGGG | Predicted or validated ATF4 target |
| **VEGFA.g4** | VEGFA | CTGCCATCCAATCGAGACCCTGG | Predicted or validated ATF4 target |
| **VEGFA.g5** | VEGFA | CCATCCAATCGAGACCCTGGTGG | Predicted or validated ATF4 target |
