## Supplementary material for "Mitochondrial Integrated Stress Response Activation Creates a Therapeutic Vulnerability to MCL-1 Inhibition in Acute Myeloid Leukemia": Sup Table 2

**Table S2: Primer and probe set sequences used to determine viral titer.**

| **Target** | **Primer** | **Probe** |
| --- | --- | --- |
| **HIV Psi** | 5’-ACTTGAAAGCGAAAGGGAAAC-3’  5’ -CAC CCATCTCTCTCCTTCTAGCC- 3’ | 5’-56-FAM-AGCTCTCTC-ZEN-GACGCAGGACTCGGC-3IABkFQ-30 |
| **RPP30** | 5’ -GCGGCT GTCTCCACAAGT-3’  5’ -GATTTGGACCTGCGAGCG-3’ | 5’-5HEX-CTGACCTGA-ZEN-AGGCTCT-3IABkFQ-3’ |
