## Supplementary material for "Mitochondrial Integrated Stress Response Activation Creates a Therapeutic Vulnerability to MCL-1 Inhibition in Acute Myeloid Leukemia": Sup Table 3

**Table S3: CRISPR-Cas9 editing construct sequences.** Guides used to target the genes presented in this study. Primer sequences used for NGS to detect gene INDELS.

| **Name** | **Sequence (5’ to 3’)** |
| --- | --- |
| **sgRNA spacer sequence** | |
| CAGE2466.ATF4.g1 | UCAGAACAGCUAACCUCUAA |
| CAGE2470.ATF4.g4 | UGGGUGGGUGGCGCUUCACU |
| CAGE2468.DELE1.g2 | ACUGGGACCUAGCCUCUGGA |
| CAGE410.EIF2AK1.g4 | UUGUUGGCUAUCACACCGCG |
| CAGE237.BAK.g2 | GCUCACCUGCUAGGUUGCAG |
| CAGE357.BAX.g1 | CUGCAGGAUGAUUGCCGCCG |
| **NGS primer sequences (NGS overhangs in upper case)** | |
| CAGE2466.ATF4.DS.F | CTACACGACGCTCTTCCGATCTcccctggtctccgtgagcgtccatt |
| CAGE2470.ATF4.DS.R | CAGACGTGTGCTCTTCCGATCTgccaactatacggctccagggcc |
| CAGE2468.DELE1.DS.F | CTACACGACGCTCTTCCGATCTacgtggatgctctaccctgagaaatc |
| CAGE2468.DELE1.DS.R | CAGACGTGTGCTCTTCCGATCTtgggcgacagaatgagactccatct |
| CAGE410.EIF2AK1.DS.F | CTACACGACGCTCTTCCGATCTaggggccatctgtattttgctctgga |
| CAGE410.EIF2AK1.DS.R | CAGACGTGTGCTCTTCCGATCTaatggtgtgaacccaggaggcggag |
| CAGE237.BAK.DS.F | CTACACGACGCTCTTCCGATCTtgactcccagctttgatcctcagagg |
| CAGE237.BAK.DS.R | CAGACGTGTGCTCTTCCGATCTgagcccaacataaaagaggagcaacc |
| CAGE357.BAX.DS.F | CTACACGACGCTCTTCCGATCTttcccttgtcccccgttggcctgtt |
| CAGE357.BAX.DS.R | CAGACGTGTGCTCTTCCGATCTctgcagctgcccaccttgagcacc |
